# DNA uptake by cell wall-deficient bacteria reveals a putative ancient macromolecule uptake mechanism

**DOI:** 10.1101/2022.01.27.478057

**Authors:** Renée Kapteijn, Shraddha Shitut, Dennis Aschmann, Le Zhang, Marit de Beer, Deniz Daviran, Rona Roverts, Anat Akiva, Gilles P. van Wezel, Alexander Kros, Dennis Claessen

## Abstract

Horizontal gene transfer in bacteria is widely believed to occur via three main mechanisms: conjugation, transduction and transformation. These mechanisms facilitate the passage of DNA across the protective cell wall using sophisticated machinery. We present here a new mechanism of DNA uptake that is independent of canonical DNA uptake machineries and is used by bacteria that live without a cell wall. We show that the cell wall-deficient bacteria engulf extracellular material, whereby intracellular vesicles are formed, and DNA is internalized. This mechanism is not specific to DNA, and allows uptake of other macromolecules and even 125 nm lipid nanoparticles (LNPs). Uptake was prevented by molecules known to inhibit eukaryotic endocytosis, suggesting this to be an energy-dependent process. Given that cell wall-deficient bacteria are considered a model for early life forms, our work provides a possible mechanism for primordial cells to acquire new genetic material or food before invention of the bacterial cell wall.

## INTRODUCTION

Bacteria are constantly exposed to changing environmental conditions and rely on their cell envelope for protection. The cell envelope consists of a cell membrane and a cell wall to separate the internal from the external environment. The cell membrane is a phospholipid bilayer that encloses the cytoplasm and functions as a selective barrier. The cell wall consists of a thick peptidoglycan (PG) layer for Gram-positive bacteria and a thinner PG layer surrounded by an outer membrane for Gram-negative bacteria. The peptidoglycan layer is an important mesh-like structure that not only provides protection against mechanical stress and turgor pressure, but also defines cell shape and rigidity.

To facilitate the selective passage of macromolecules across the cell envelope, bacteria have evolved specialized and sophisticated transport systems (Costa et al., 2015; Forster and Marquis, 2012). For instance, naturally transformable bacteria rely on protein complexes for DNA uptake, with components similar to type IV pili or type II secretion systems. Active transport of DNA across the cell wall is facilitated by the retraction of pilus structures that bind DNA (Chen and Dubnau, 2004; Ellison et al., 2018). Alternatively, the assembly and release of short pilus structures are thought to create transient holes in the PG layer that allow DNA to diffuse to the cell membrane (Muschiol et al., 2015). DNA binding and pore-forming proteins are then used to translocate the DNA across the cell membrane.

Although the cell wall is a vital structure for most bacteria, some bacteria naturally lack a cell wall, or can shed their wall under specific conditions. Examples include the members of the Mollicutes, that are parasitic and live in specific osmotically protective environments such as human mucosal surfaces or the phloem sieve tubes of plants (Stülke et al., 2009). Prolonged exposure to environmental stressors such as cell wall-targeting agents generates so-called L-forms, which are cells that proliferate without their cell wall. Reproduction of L-forms is driven by the upregulation of membrane synthesis and is characterized by blebbing, tubulation and vesiculation (Mercier et al., 2013; Ramijan et al., 2018). These primitive cell-like characteristics make L-forms an attractive model system to study the evolution of early life (Briers et al., 2012b; Errington et al., 2016). How the absence of the cell wall affects uptake of macromolecules such as DNA from the environment is unknown.

In this study we show that L-forms of the filamentous actinomycete *Kitasatospora viridifaciens* can naturally take up DNA independent of the canonical DNA translocation machinery. Instead, uptake is facilitated by a new mechanism of horizontal gene transfer that involves the invagination of the cell membrane leading to internal vesicle formation. Furthermore we show that this mechanism is robust and allows the non-specific uptake of other macromolecules from the environment as well. Given that L-forms are considered a model for early cellular life, our work provides insight into how such ancient cells may have acquired large biomolecules and nanoparticles from the environment without the need for complex transport machineries.

## RESULTS

### Natural and artificial DNA uptake by wall-deficient cells

*K. viridifacien*s is a mycelium-forming bacterium that can extrude temporary wall-deficient cells, called S-cells, under conditions of osmotic stress (Ramijan et al., 2018). These cells can only proliferate by rebuilding the cell wall and reverting to a mycelial mode-of-growth, similar to artificially created cell-wall deficient protoplasts. Prolonged incubation of S-cells in high osmotic pressure can induce the switch to an L-form state that allows reproduction without rebuilding the cell wall. It is unknown whether these cell wall-deficient cells can take up DNA via natural transformation. To analyse this, protoplasts, S-cells and L-forms (*alpha*) were incubated with plasmid DNA, and subsequently plated on selective and non-selective medium (Figure 1A). Notably, L-forms were consistently able to take up DNA, unlike protoplasts or S-cells (Figure 1B). This DNA uptake ability was not restricted to the penicillin-induced L-forms (lines *alpha* and *delta*), as the osmotically induced L-form (line M1) (Ramijan et al., 2018) could also take up plasmid DNA. No transformants were obtained with *alpha* when intact or fragmented genomic DNA was used (Figure S1A). While natural transformation was restricted to L-forms, all wall-deficient cells could be chemically transformed using polyethylene glycol (PEG), with protoplasts, S-cells and L-forms having an average transformation efficiency between 1.7 – 2.5% (Figure S1B). The addition of PEG also enabled transformation of *alpha* with genomic DNA, even if this was present in a crude cell extract. On the other hand, use of methylated DNA prevented transformation, indicating that transformation is possible with different types of DNA, but can be limited by the presence of a different methylation pattern (Figure S1C). By contrast, walled cells could not be transformed either with or without PEG (Figure 1B and Figure S1B). These results show that proliferating wall-deficient L-forms can take up DNA naturally, while walled cells and transient wall-deficient S-cells and protoplasts cannot.

**Figure 1.**
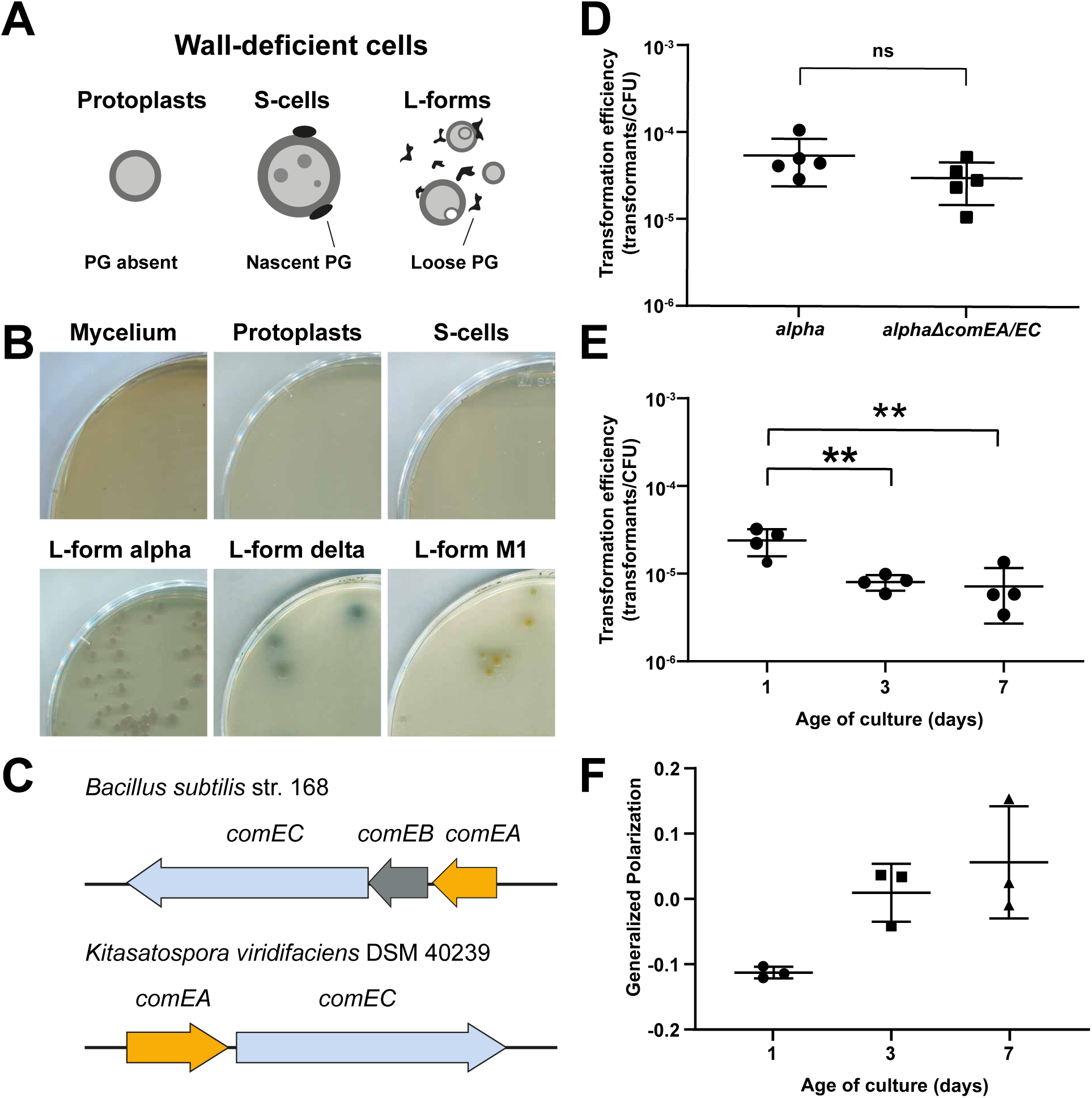
Natural DNA Uptake of Wall-Deficient Cells Is Independent of the Competence Proteins ComEA and ComEC and Correlates with Membrane Fluidity. (A) Schematic representation of the different wall-deficient cell types of *K. viridifaciens* that can be created artificially (protoplasts) or naturally (S-cells and L-forms). PG = peptidoglycan. (B) Mycelium, protoplasts, S-cells and L-form lines *alpha*, M1 and *delta* were incubated with plasmid DNA (pRed*) for 24h, plated on selective medium and incubated at 30°C to select for transformed cells. Note that only L-forms show consistent DNA uptake. (C) Localization of putative ComEA and ComEC genes (BOQ63_29625 and BOQ63_29630, respectively) on the chromosome of *K. viridifaciens* DSM 40239 as compared to *comEC* and *comEA* of naturally transformable *Bacillus subtilis* str. 168. (D) Natural transformation assay of 7-day of alpha and *alphaΔcomEA/EC* using pFL-*ssgB*. ns = not significant (n=5 replicates, two-tailed independent t-test, t(8)=1.572, P=0.155). Data are represented as mean ±SD with individual data points. (E) Natural transformation efficiency of 1-, 3- and 7-day old alpha after 24 h incubation with pFL-ssgB. Asterisks indicate statistically significant different transformation efficiency (n=4 replicates, one-way ANOVA, F (2,9) = 12.16, Tukey post-hoc test, P =.006 (1-3 day) and .005 (1-7 day)). Data are represented as mean ±SD with individual data points. (F) Generalized polarization as measurement of membrane fluidity of 1-, 3- and 7-day old alpha as calculated from the shift in the fluorescence emission spectrum of the membrane dye Laurdan. Lower GP indicates a higher membrane fluidity. Data are represented as mean ±SD with individual data points, n=3.

### L-forms take up DNA in the absence of canonical DNA translocation machinery

Naturally transformable bacteria use a specialized DNA translocation machinery with similarities to type IV pili or type II secretion systems to take up external DNA (Chen and Dubnau, 2004). Similar components of this canonical system might also be involved in DNA uptake by L-forms. A BlastP search using the DNA-binding protein ComEA and channel protein ComEC of the naturally transformable bacterium *Bacillus subtilis* against *K. viridifaciens* yielded two significant hits: BOQ63_29625 (helix-hairpin-helix domain-containing protein) and BOQ63_29630 (ComEC/Rec2 family competence protein), respectively (Figure 1C and Table S1). The *B. subtilis* helicase/DNA translocase ComFA resulted in a hit to a putative Mfd-encoding gene (BOQ63_20315), a widely conserved bacterial protein that mediates transcription-coupled DNA repair (Roberts and Park, 2004). No other orthologues were found that correlated to proteins involved in DNA transport across the cell envelope for *B. subtilis*, the Gram-negative *Neisseria gonorrhoeae* (Kruger and Stingl, 2011) or for the T4SS-related DNA uptake system of *Helicobacter pylori* (Gilbreath et al., 2011) (Table S1). L-forms lack an intact peptidoglycan-based cell wall and therefore DNA must only cross the cell membrane for internalization. As ComEA and ComEC function in DNA transport across the cell membrane (Friedrich et al., 2001; Inamine and Dubnau, 1995; Kruger and Stingl, 2011) we wondered whether these proteins are involved in DNA uptake in L-forms. Therefore, we replaced the putative *comEC* and *comEA* genes in the L-form strain *alpha* by an apramycin resistance cassette (Figure S1D). Strikingly, the simultaneous deletion of the *comEA* and *comEC* genes did not affect the natural transformation efficiency (two-tailed independent t-test, *t*(8)=1.572, *P*=0.155), indicating that DNA uptake by L-forms occurs independent of this canonical DNA translocation machinery (Figure 1D).

### High membrane fluidity is not sufficient for natural DNA uptake in wall-deficient cells

One of the factors controlling the development of competence for DNA uptake in *B. subtilis* is the growth phase (Dubnau, 1991; Hamoen et al., 2003). To study if culture age is also affecting the DNA uptake ability of L-forms, differently aged cultures were subjected to a natural transformation assay. One-day old cultures of *alpha* take up DNA more easily than 3- or 7-day old cultures (one-way ANOVA, F (2,9) = 12.16, Tukey post-hoc test, *P* =.006 and .005 respectively) (Figure 1E). It is not unlikely that differences in membrane properties that occur during cellular growth may in turn affect the DNA uptake ability. Membrane fluidity is a measure for the average viscosity of the lipid bilayer, which can affect the positioning and movement of proteins and lipids within the membrane (Lenaz, 1987). A higher membrane fluidity is characterized by increased fatty acid disorder, lower lipid packing and higher diffusion rates, which can lead to increased membrane permeabilization (Chapman, 1975; Lande et al., 1995). Analysis of the membrane fluidity of the differently aged cultures indicated that the increased DNA uptake ability may correlate positively with the fluidity of the membrane, as deduced from the generalized polarization (GP) (Scheinpflug et al., 2017) (Figure 1F), although no statistical significant differences were observed (Welch ANOVA, F(2, 2.798), with Games-Howell post-hoc test: 1-3 day *P* = .068; 1-7 day *P*= 0.134; 3-7 day *P* = 0.711). A relatively low fluidity might explain why temporary wall-deficient protoplasts and S-cells cannot take up DNA naturally. However, the fluidity of protoplasts was within the range of 1- to 7-day-old cultures as measured using a plate assay (Figure S1E). Subsequent analysis of the GP by fluorescence microscopy imaging showed that although protoplasts and S-cells tend to have less fluid membranes, these values stay within the range of the membrane fluidity of 1- to 7-day old L-forms (Figure S1F). Therefore, although membrane fluidity may contribute to efficient DNA uptake, it is not sufficient to explain this process.

### L-forms take up DNA via an endocytosis-like mechanism

To further investigate the mechanism facilitating DNA uptake by L-forms, we added Cy5-labelled plasmid DNA to L-forms expressing cytosolic eGFP. Labelled plasmid DNA was found either on the outside of the L-form cell membrane, or within apparent internal vesicles (Figure 2A and control Figure S2A). As these internal vesicles were devoid of eGFP, we reasoned that they could have originated by an invagination process of the membrane, whereby extracellular material becomes trapped inside the vesicles. To test this directly, we incubated eGFP-expressing L-forms with the fluorescent dye SynapseRed C2M (SynapseRed). Given that SynapseRed cannot diffuse through the cell membrane, any fluorescent signal on the membranes surrounding internal vesicles would be a strong argument that such vesicles were derived from the cell membrane. Indeed, SynapseRed was found to not only stain the cell membrane of the L-forms but also the membranes of internal vesicles after overnight incubation (Figure 2B). Staining with SYTO-9 further indicated that chromosomal DNA was present in the cytosol but not inside internal vesicles (Figure S2B). Incubation of protoplasts producing cytosolic eGFP with Synapse Red showed that areas with less cytosolic fluorescence emission were caused by internal membrane structures rather than by formation of internal vesicles (Figure S2C). Similar incubation of S-cells showed the presence of internal vesicle-like structures. However, unlike for L-forms, subsequent staining of S-cells of a strain producing cytosolic-mCherry with SYTO-9 indicated that these vesicles were filled with chromosomal DNA. This indicates that internal structures observed in protoplasts and S-cells are not the same internal vesicles as those seen in L-forms and may not be involved uptake of external fluids. Taken together, these results strongly suggest that the observed vesicles inside L-forms originate from invagination of the cell membrane whereby extracellular material may become trapped inside such vesicles.

**Figure 2.**
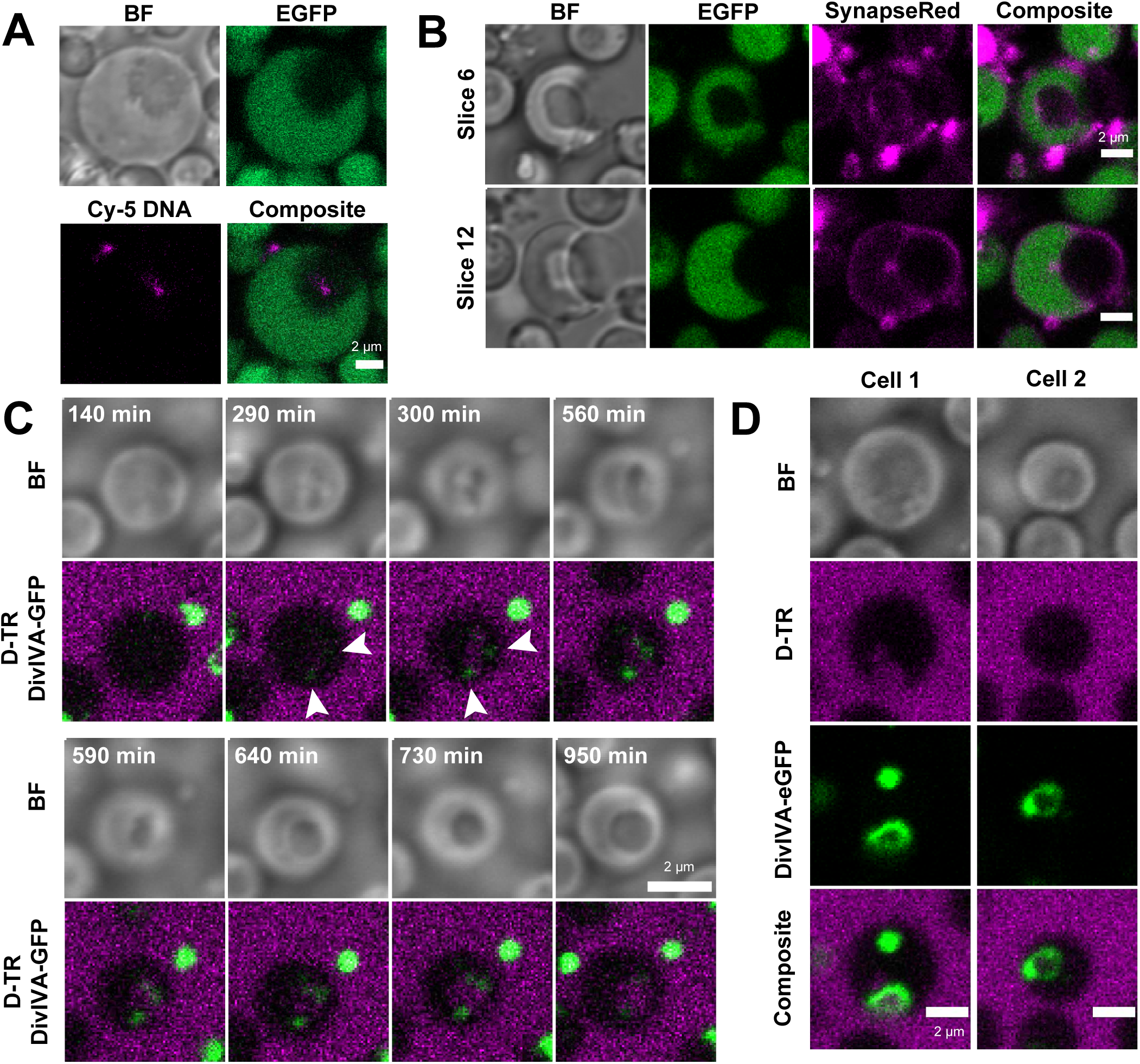
Formation of Internal Vesicles and Uptake of External Fluids in L-forms. (A) Fluorescence micrograph of *alpha* pIJ82-GFP (cytoplasmic eGFP; green) incubated with Cy-5 labelled plasmid DNA (pFL-*ssgB*; magenta). BF = Brightfield. Scale bar = 2 μm. (B) Incubation of *alpha* pIJ82-GFP with the membrane-impermeable dye SynapseRed C2M (SynapseRed; magenta), showing two z-slices of one L-form cell. BF = brightfield. Scale bar = 2μm. (C) Stills of a time-lapse imaging experiment of *alpha* producing DivIVA-eGFP (*alpha* pKR2) (green) incubated with 3 kDa Dextran-Texas Red (D-TR; magenta). Arrows indicate localization of DivIVA-eGFP. Scale bar = 2μm. See also Video S1. (D) Formation of foci and ring-structures of DivIVA-eGFP in *alpha* pKR2 (green) incubated with Dextran-Texas Red (D-TR, magenta). Scale bar = 2μm. Note that L-forms are able to take up fluorescently stained DNA and Dextran by formation of internal vesicles.

In eukaryotes, endocytosis is a process that enables the uptake of external cargo via internal vesicle formation, which is eventually degraded or recycled (Cossart and Helenius, 2014; Elkin et al., 2016). Fluorescently labelled dextrans are widely used as markers for endocytosis in eukaryotes (Araki et al., 1996; Li et al., 2015). To identify if such an endocytosis-like process could be present in L-forms and to visualize the uptake of external materials, we incubated the cells with Dextran Texas-Red (D-TR) and performed time-lapse imaging. The L-form strain used also expresses DivIVA-eGFP, which has strong affinity for negatively curved membrane regions (*alpha* pKR2) (Hammond et al., 2019; Jurasek et al., 2020). Such regions are expected to be formed upon invagination of the membrane. After 290 minutes of incubation, D-TR was visible inside the L-form and faint spots of DivIVA-eGFP started to appear adjacent to this region (Figure 2C and Video S1). This progressed to a clear inward bulging of the cell membrane with two foci of DivIVA-eGFP on either side of the invaginated membrane and an inflow of D-TR (t=560 min). After 640 min an internal vesicle was formed that contained D-TR. In other cells, DivIVA-eGFP appeared to form a ring-like structure, which sometimes enveloped the invaginating membrane (Figure 2D cell 1 and 2 respectively). The presence of DivIVA near the site of invagination implies the presence of negatively curved regions in the membrane. Notably, DivIVA is not required for vesicle formation or DNA uptake, as the deletion of *divIVA* in *alpha* (*alpha*ΔDivIVA) had no effect on natural transformation (two-tailed independent t-test, *t*(8)=0.489, *P*=0.638) (Figure S2D), and internal vesicles were still formed by this strain (Figure S2E). Furthermore, internalization of D-TR was also observed in L-forms that did not express DivIVA-eGFP, indicating that uptake is not a consequence of the presence of the fusion protein (Figure S2F). Incubation of protoplasts and S-cells with D-TR up to 72 h did not result in D-TR encapsulation in internal vesicles (Figure S2G). Altogether, these results show that the invagination of the cell membrane of L-forms can lead to internal vesicle formation and may represent an endocytosis-like mechanism allowing uptake of molecules, including DNA, from the environment.

### Lipid nanoparticles are internalized in vesicles in an energy-dependent manner

Lipid nanoparticles (LNPs) are non-viral particles that are used to deliver nucleic acids and drugs to human cells via endocytosis (Cullis and Hope, 2017). LNPs do not have a lipid bilayer structure, but consist of an electron-dense, hydrophobic core of lipids that encapsulate nucleic acids by electrostatic interactions and are surrounded by a layer of PEG-lipids (Cullis and Hope, 2017; Evers et al., 2018; Hou et al., 2021). Once the endosome acidifies the ionizable lipids become positively charged, which allows the LNP to destabilize the endosome membrane and deliver its cargo into the cell. LNPs can also be fluorescently tagged by the incorporation of fluorophore-conjugated phospholipids (Kulkarni et al., 2019). To further explore the ability of L-forms to take up external particles, the cells were incubated with rhodamine-labelled LNPs (LNP-LR, containing 18:1 Liss Rhod PE) with an average size of 125 nm to allow their detection inside L-forms. After addition of LNP-LR to 7-day-old L-forms, clear foci could be detected inside the cells after overnight incubation, as well as localization of LNPs to the cell membrane (Figure 3A and Figure S3B and S3C). When L-forms were used that express eGFP in the cytosol, vesicles only contained LNPs and not eGFP, strongly suggesting that the LNPs had been internalized in vesicles devoid of the cytoplasm (Figure 3B, C). Importantly, internalization of LNP-LR by L-forms could be blocked by the addition of sodium azide (1, 2.5 and 10 mM) or incubation of cells at 4°C, conditions that are commonly used to inhibit endocytosis (Atkinson et al., 2002; Hoffmann and Mendgen, 1998; Sato et al., 2009; Subramanya et al., 2009). Under such conditions, the LNPs only localized to the cell membrane rather than forming foci inside the cell (Figure 3D, E and Figure S3D-E). These results are consistent with an uptake process of LNPs that is energy-dependent, whereby the particles are internalized by a membrane invagination process.

**Figure 3.**
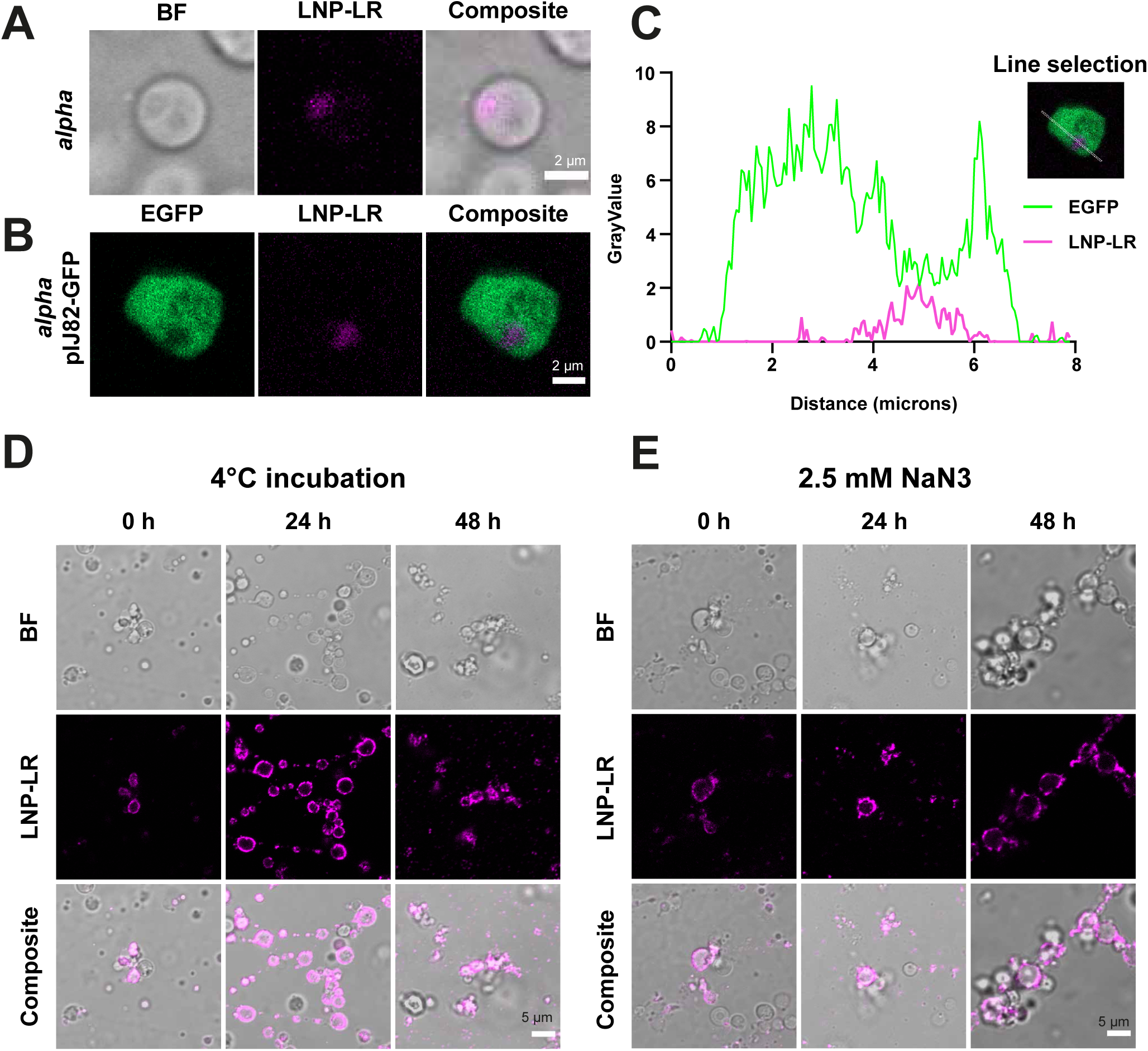
Localization of Lipid Nanoparticles in Internal L-form Vesicles. (A-B) Localization of LNP-LR (Lipid Nanoparticle containing 18:1 Liss Rhod PE; magenta) in internal vesicles of *alpha* (A) and *alpha* pIJ82-GFP (B) after overnight or 3-day incubation at 30°C respectively. Scale bar = 2μm. (C) Density profile plot and corresponding line selection of *alpha* pIJ82-GFP incubated with LNP-LR showing a decrease in cytoplasmic eGFP emission correlates with an increase in LNP-LR emission. (D-E) Localization of LNP-LR during incubation with *alpha* at 4°C (D) or in the presence of 2.5 mM sodium azide at 30°C (E) after 0, 24 and 48 h incubation. Similar results were obtained with 1 and 10 mM sodium azide (data not shown). Scale bar = 5 μm. Note that incubation of L-forms with lipid nanoparticles (average size of 125 nm) results in their localization inside internal vesicles, a process that can be inhibited by incubation at 4°C or sodium azide.

### High-resolution imaging of L-forms using cryo-FIB-SEM

To better understand their ultrastructure and composition, the intracellular vesicles were imaged using 3D cryo-correlative light and electron microscopy (cryo-CLEM) (Figure 4A). Cryo-FIB-SEM (Focused Ion Beam - Scanning Electron Microscopy) allows the 3D high resolution imaging of L-forms and internal vesicles. The cryogenic sample preparation and imaging ensures that the L-forms are visualized in a near-to-native state (Shimoni and Muller, 1998; Studer et al., 1989).

**Figure 4.**
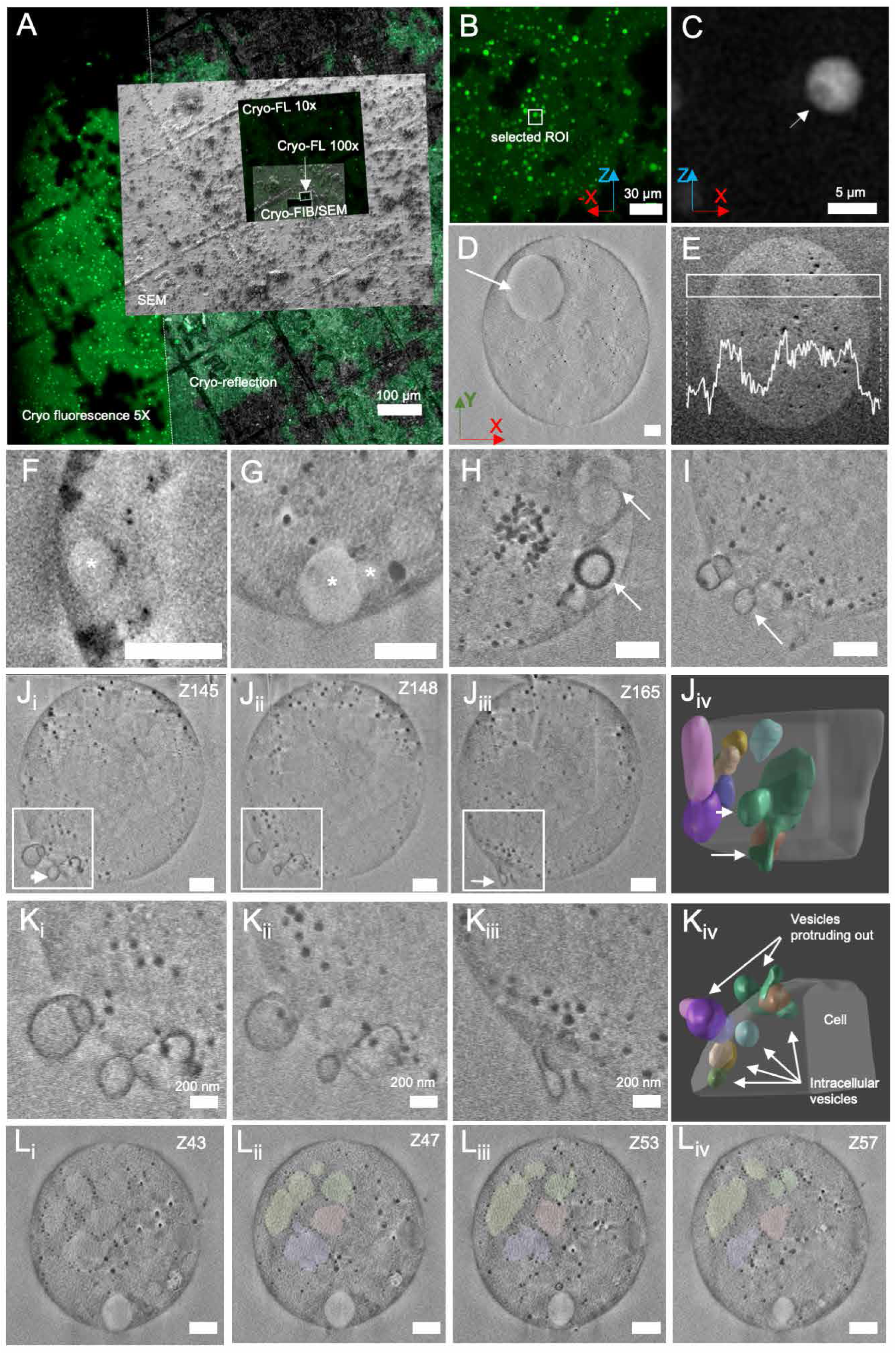
3D Cryo-Fluorescence and Cryo-FIB-SEM of L-forms Reveals its Ultra-Structure in High Resolution. (A) Correlated fluorescence and electron micrographs of the frozen sample (Zen Connect image). The bright green dots indicate individual cells of *alpha* pIJ82-GFP. A finderTOP raster visible both in fluorescence and electron microscopy facilitates alignment between the two imaging modules. The small squares indicate different regions of interest, imaged at higher resolution. FL: Fluorescence light (B) Higher resolution image of one region of interest, showing many fluorescent cells. (C) L-form depicted by white box in B, showing intracellular dark sphere (∼ 1 micrometer, white arrow). (D) SEM image (SE, Inlens) of cell in C) with white arrow indicating the internal vesicle. The X, Y and Z arrows in B, C and D indicate the 3D orientation of the imaged cell as observed in 3D FIB-SEM. (E) Superposition of five consecutive slices (backscattered images) of cell in D). Inset: Intensity plot profile (white) of the region in white box. (F-I) FIB-SEM slices showing different types of internal vesicles. (F-G) Vesicles lining the cell membrane. Asterisks indicate vesicles. (H) Vesicle complex, note the different membrane thickness of vesicles indicated with white arrows. See also Figure S6D and Video S3. (I) Membrane protrusions as indicated with white arrow. (J-K) Analysis of the interconnected vesicles of the cell in I). (Ji-iii) Three consecutive slices showing the interaction of different vesicles. Ki-iii show higher magnification of the regions in white boxes in Ji-iii, respectively). (Jiv, Kiv) 3D segmentation of Ki-iii. While some of the vesicles are intracellular, others protrude out of the cell. A complete connected vesicle structure is shown in green and is indicated by white arrows in I, Jiii and Jiv. See also Figure S6A-C and Video S2. (L) Regions with different contrast are lined with black particles representing putative lipid bodies. The size distribution of the black particles is between 25 to 60 nm. Scale bars represent 500 nm unless otherwise specified.

Following high-pressure freezing, cells with putative intracellular vesicles were detected based on internal darker regions lacking cytosolic eGFP using *alpha* pIJ82-GFP (Figure S4). Specific L-forms (example of selection in Figure 4B, C) were imaged in detail using cryo-FIB-SEM. The reduction in cytosolic eGFP indeed matched the presence of internal vesicles as detected by FIB-SEM (Figure 4C, D, white arrow), in line with previous results (Figure 2B). In addition, the composition of the cytoplasm and internal vesicle content was different, as measured using the InLens energy selective backscattered (EsB) detector which provides contrast based on the distribution of heavier elements (Figure 4E). Analysis of the pixel intensity indicated that the contrast level inside the internal vesicle was similar to the extracellular environment, whereas the cytoplasm had a higher contrast. Moreover, an over-exposure experiment showed that the vesicle has the same capacity to absorb the electron dose as the medium outside, different from the rest of the cell (Figure S5A-B). These results support the finding that internal vesicles contain extracellular medium and are formed via membrane invagination (Figure 2C).

Further high-resolution imaging indicated the presence of multiple internal vesicles within individual cells (Figure 4F-I, Figure S5C-E). Most detected vesicles were lining the cell membrane (Figure 4G, Figure S5C-E), varied in size and membrane thickness (Figure 4H) and could even be present inside larger vesicles (Figure 4H and Figure S6D), like the previously observed secondary vesicle (Figure S2F). In addition, vesicles could be observed budding out of the cell membrane (Figure 4I). 3D reconstruction of the budding vesicles based on contour tracing revealed that these were either an extension of an internal vesicle, or remained connected to internal vesicles, forming a complex (Figure 4J-K, Figure S6A-D, Video S2 and S3).

In some cases, cells contained intracellular regions with different grey values from the rest of the cell (Figure 4Li). These regions had a size distribution of 300 to 800 nm, did not line the cell membrane, and were surrounded by dark particles of around 25-60 nm in diameter (Figure 4Lii-iv). It could be possible that these dark particles are lipid bodies, compared to previous cryo-FIB-SEM observations (Spehner et al., 2020; Vidavsky et al., 2016). A potential interpretation is that the internal regions are vesicles of which the enclosing lipid membrane has partially degraded. The lipids and lipidic degradation products may have accumulated in lipid droplets that result in the observed black particles.

These results further confirm that the internal vesicles observed in *K. viridifaciens* L-forms contain external medium and can be formed by invagination of the cell membrane. L-forms can contain multiple vesicles of varying sizes, in some cases forming clusters or complexes of vesicles that can protrude out of the cell membrane. Internal vesicles may release their contents in the cell after vesicle degradation. These findings support a model for uptake of macromolecules such as DNA by engulfment, followed by release of the cargo after vesicle disruption (Figure 5).

**Figure 5.**
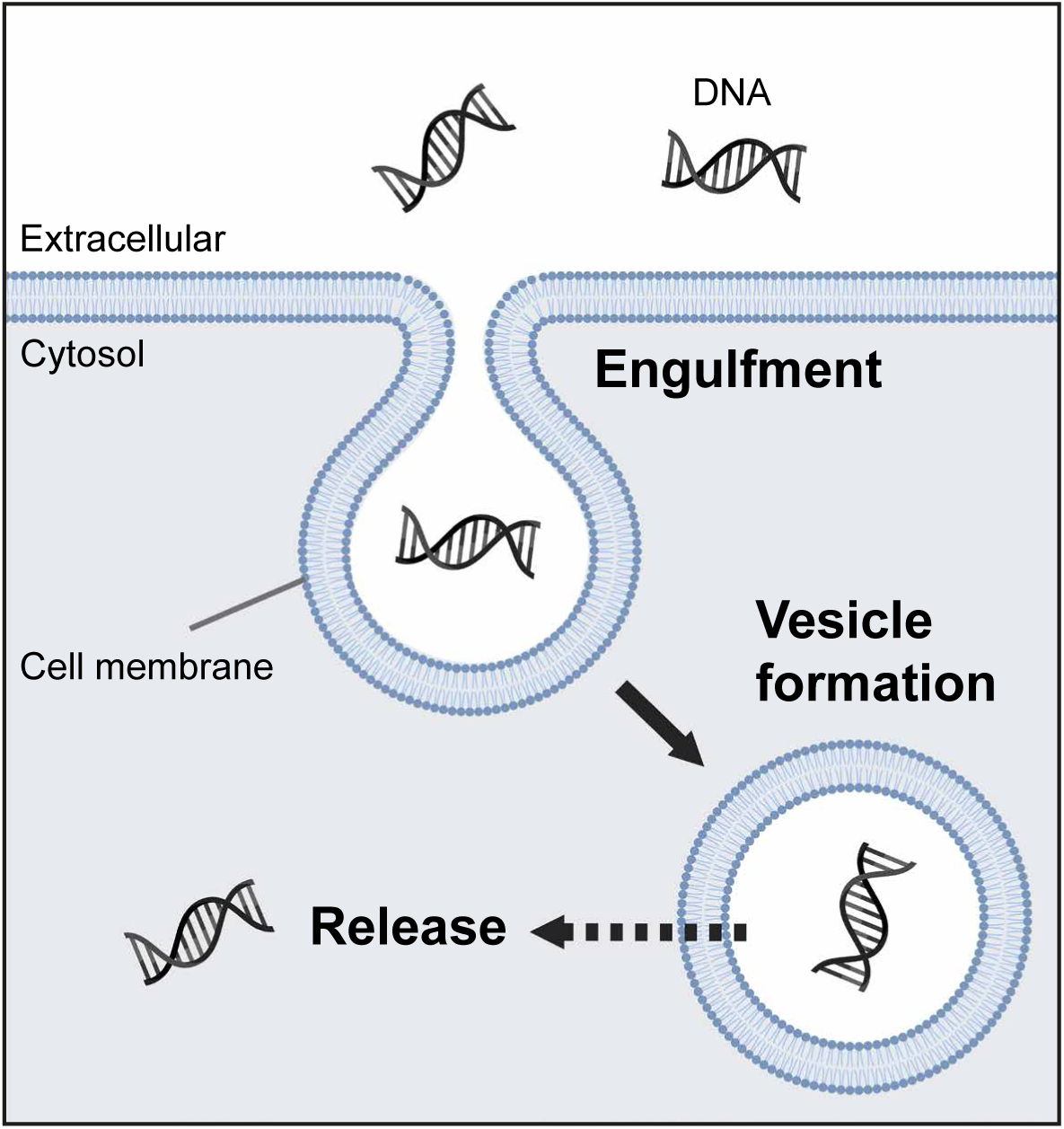
Proposed Model for DNA Uptake by Internal Vesicle Formation in L-forms. Excess membrane synthesis results in invagination of the cell membrane, leading to the formation of internal vesicles in L-forms. In this process, extracellular liquid containing DNA or other macromolecules is engulfed. Finally, DNA is released from internal vesicles by an unknown process (indicated by dashed arrow), which may involve vesicle disruption. Image created with BioRender.com.

## DISCUSSION

The bacterial cell wall is an important protective barrier towards the environment, providing stress resistance and enabling the selective passage of molecules. However, in recent years it has become clear that under some conditions, bacteria may also thrive without this layer. Prolonged exposure to environmental stresses, such as cell-wall targeting agents or a high osmotic pressure, can induce the formation of L-forms that efficiently proliferate without their cell wall (Allan et al., 2009; Ramijan et al., 2018). The consequences of such a wall-deficient bacterial lifestyle on their ability to take up DNA are largely unknown. Here we provide evidence that L-forms may take up DNA and other macromolecules via engulfment and the subsequent formation of internal vesicles (Figure 5).

### A new mechanism for HGT?

Well-known mechanisms for HGT are natural transformation, transduction, and conjugation (as reviewed in Arnold et al., 2021; Thomas and Nielsen, 2005). These mechanisms require sophisticated machinery to enable transport of DNA across the cell envelope. We here show that wall-deficient cells such as protoplasts, S-cells and L-forms of *K. viridifaciens* take up DNA using PEG. Importantly, L-forms are the only wall-deficient cells that achieve natural transformation using plasmid DNA without PEG. Naturally transformable bacteria use a canonical and complex system for DNA uptake across the cell wall and cell membrane. The latter step requires the DNA-binding protein ComEA and the pore-forming channel protein ComEC, with homologs found across naturally transformable Gram-positive and Gram-negative species (e.g., ComE and ComA in *N. gonorrhoeae*). Disruption of either of these proteins typically results in a drastic reduction or even absence of transformation (Friedrich et al., 2001; Hahn et al., 1987; Inamine and Dubnau, 1995; Yeh et al., 2003). However, disruption of the likely genes for ComEA and ComEC in L-forms of *K. viridifaciens* had no effect on the ability to take up DNA, suggesting a mechanism independent of the canonical DNA translocation machinery.

### An endocytosis-like process in L-form bacteria

Endocytosis is a fundamental and highly regulated process in eukaryotes that is involved in the uptake of nutrients, regulation of plasma membrane composition, sensing of the extracellular environment and signaling (Thottacherry et al., 2019). Invagination of the membrane and subsequent membrane scission and vesicle formation allows cells to internalize a wide array of cargo such as fluids, ligands, plasma membrane proteins and sometimes even entire bacteria. Invagination is often followed by passing the cargo through the endosomal pathway and lysosomal degradation (Cossart and Helenius, 2014). Specific mammalian cells can take up DNA, followed by active gene expression (Wolff et al., 1990), which potentially occurs via endocytosis, although the exact mechanism is unclear (reviewed by Budker et al., 2000; Trombone et al., 2007; Wolff and Budker, 2005).

This work shows that L-forms use an endocytosis-like mechanism for the uptake of DNA, whereby membrane invagination led to the formation of intracellular vesicles that during their formation encapsulated extracellular material (Figure 5). Via this process, not only DNA but also other macromolecules such as 3 kDa dextran and even 125-nm lipid nanoparticles were taken up, strongly suggesting that the uptake process is non-specific. Interestingly, an older study also reports the uptake of fluorescent dextrans in internal vesicles of *Bacillus subtilis* L-forms, which was proposed to occur via fluid-phase endocytosis (Oparka et al., 1993).

The mechanism underlying formation of intracellular vesicles in L-forms most likely depends on increased membrane dynamics due to excess membrane synthesis (Mercier et al., 2013; Studer et al., 2016). An imbalance in the cell surface to volume ratio due to excess membrane synthesis can lead to internal vesicle formation in spherical *E. coli* and *B. subtilis* shape mutants (Bendezu and de Boer, 2008; Mercier et al., 2013). Internal vesicles or vacuoles can also be formed in enlarged protoplasts and spheroplasts (containing an outer membrane) which are maintained in conditions that allow cell membrane expansion (Nishida, 2020; Takahashi et al., 2020). Indeed, a lack of excess membrane production may also explain why we did not observe consistent DNA uptake in protoplasts and S-cells, both of which are unable to proliferate without their wall.

High-resolution electron microscopy imaging revealed multiple internal vesicles inside L-forms. Interestingly, the L-forms also contained regions not surrounded by a membrane but were lined with darker spots that may represent lipid bodies (Spehner et al., 2020; Vidavsky et al., 2016), possibly originating from the degradation products of the membrane of internal vesicles. This disintegration would lead to release of the cargo into the cytoplasm. In eukaryotes, escape of therapeutics from endosomal vesicles can be mediated by bacterial, viral, and chemical agents or by nanoparticles (Patel et al., 2019; Varkouhi et al., 2011). Escape mechanisms include pore formation, destabilization of the membrane, nanoparticle swelling or osmotic rupture. High sucrose levels or the proton sponge effect facilitate the influx of protons followed by chloride ion accumulation and inflow of water, leading to rupture of the vesicle (Behr, 1997; Cervia et al., 2017; Ciftci and Levy, 2001; Liang and W. Lam, 2012). Acidification of endosomes occurs via membrane-localized vacuolar ATPases (V-ATPases) that pump protons into the vesicles (Forgac, 2007). Bacteria have similar proton pumps called F-ATPases on their plasma membrane and have been found on the membrane of intracellular vesicles of enlarged protoplasts (Hensel et al., 1996; Mulkidjanian et al., 2007; Takahashi et al., 2020). Considering the complexity of known escape mechanisms further research is required to understand if and how internal L-form vesicles can disintegrate to release their contents in the cytoplasm.

### An endocytosis-like mechanism for macromolecule uptake in primordial cells

L-forms have been proposed as a model to study early lifeforms due to their lack of cell wall and biophysical way of proliferation (Briers et al., 2012b; Errington et al., 2016). Horizontal gene transfer is thought to have played a pivotal role in the evolution of early life (Woese, 1998; Woese, 2000). This may have occurred in cells that did not yet evolve a cell wall, allowing genetic recombination after cell fusion or lightning-triggered electroporation (Errington, 2013; Kotnik, 2013), yet other mechanisms of HGT were unknown. Internal vesicles have also been observed in L-forms of other species, with varying functions and mechanisms of vesicle formation described (Han et al., 2015; Yabu, 1991) . L-forms of *Listeria monocytogenes* are capable of forming DNA-containing internal vesicles along the inside of the cell, which upon release become metabolically active (Briers et al., 2012a; Dell’Era et al., 2009), as well as forming internal vesicles via membrane invagination (Studer et al., 2016). Additionally, secondary invagination of the vesicle membrane itself can result in vesicles containing cytoplasm and represent viable offspring.

These examples provide additional support for the existence of bacterial endocytosis, and we therefore propose that this may reflect an ancient mechanism that has been retained in modern cells to allow shedding their cell wall when the environmental conditions require it. These examples provide additional support for the existence of bacterial endocytosis, and we therefore propose that this may reflect an ancient mechanism of how primordial cells acquired new genetic material and nutrients via engulfment.

In conclusion, our work shows that the permanent loss of the bacterial cell wall allows the uptake of DNA, dextran and 125 nm-sized lipid nanoparticles via internal vesicle formation. The invagination of the cell membrane, likely driven by excess membrane production, leads to the engulfment of external fluids and subsequent vesicle formation. This is an energy-dependent process that has similarities to a simple form of endocytosis as seen in eukaryotes. Future studies are required to further understand the molecular mechanisms behind this process.

## AUTHOR CONTRIBUTIONS

R.K. and S.S. carried out the experiments. L.Z., S.S. and R.K. created the plasmids and D.A provided the lipid nanoparticles. A.A., M.d.B, R.R. and D.D. performed the FIB-SEM imaging and analysis. All authors contributed to the design of the experiments and discussion of the results. R.K, D.C. and A.A. wrote the manuscript with input from all authors.

## MATERIALS AND METHODS

### Bacterial strains and culture conditions

The bacterial strains and plasmids used in this study are listed in Table S2 and S3 respectively. *Kitasatospora viridifaciens* DSM40239 (Ramijan et al., 2017) was grown confluently on maltose-yeast extract medium (MYM) to obtain spores, which were harvested after 3-4 days of growth (Stuttard, 1982). For mycelial growth in liquid, strains were grown at a density of 1 x 10^6^ spores ml^-1^ for two days in LPB medium without sucrose at 100 rpm, while LPB with sucrose was used to induce the formation of S-cells (Ramijan et al., 2018). L-forms were grown on solid L-phase medium agar (LPMA) or liquid LPB medium (Ramijan et al., 2018). Liquid cultures were inoculated with spores for *K. viridifaciens* strains or with a frozen aliquot of a 1-2-day old L-form culture in case of L-form strains. L-forms were grown in liquid culture for 3-4 days for chemical transformation and 7 days for all other experiments unless stated specifically. L-forms were adjusted to 5-7.5 x 10^7^ CFU ml^-1^ for transformation assays (based on OD_600_ of 3 for 3- and 7-day old cells and 0.2 for 1-day old cells), and 2.5-5 x 10^7^ CFU ml^-1^ (OD_600_ of 2) for all other experiments with 7-day old cells. All *Kitasatospora* cultures were grown at 30°C.

*Escherichia coli* strains were grown on solid or liquid LB medium (while shaking at 250 rpm) at 37°C. Where necessary, antibiotics (100 µg ml^-1^ ampicillin, 25 µg ml^-1^ chloramphenicol, 5 µg ml^-1^ thiostrepton, 50 µg ml^-1^ apramycin, 100 µg ml^-1^ hygromycin B with the exception of 200 µg ml^-1^ hygromycin B for LB medium) were added to the culture medium. *E. coli* JM109 (Yanisch-Perron et al., 1985) was used for cloning purposes, while *E. coli* ET12567/pUZ8002 (MacNeil et al., 1992) was used to obtain methylation-deficient DNA.

### Construction of plasmids

All PCRs were performed using PFU or Q5^®^ High-Fidelity DNA polymerase (NEB). The primers used in this study are listed in Table S4. To create pFL-*ssgB* (Table S3), a hygromycin resistance cassette was amplified using primer pair Hyg_F-231_EEV and Hyg_R+1237_HEV with pMS82 (Gregory et al., 2003) as the template. The PCR products were digested with EcoRV and cloned into pWHM3-oriT (Wu et al., 2019) to generate pWHM3-oriT-hyg (Table S3). The 3’ flank of *ssgB* was digested from pKR1 (Ramijan et al., 2018) and cloned into pWHM3-oriT-hyg using XbaI and HindIII to generate the final plasmid.

pRK1 (Table S3) was created by amplifying the upstream flanking region of *comEA* by PCR with primers FL1-comEA/comEC-FW and FL1-comEA/comEC-REV, thereby introducing unique EcoRI and XbaI resitricion sites, while the downstream flanking region of *comEC*, made by gene synthesis (Baseclear, Leiden, the Netherlands) was flanked by XbaI and HindIII sites. The flanking regions and apramycin cassette were cloned in pWHM3-oriT using the EcoRI, HindIII restriction sites interspersed with an apramycin resistance cassette containing flanking XbaI sites, thereby creating the final plasmid. The *comEA/comEC* deletion mutant was created in L-form strain *alpha* (Ramijan et al., 2018) using pRK1, which replaced the nucleotides +58 relative to the startcodon of *comEA* (BOQ63_29625) until + 2489 relative to the startcodon of *comEC* (BOQ63_29630) with an apramycin resistance cassette. Note that the gene annotation of *Streptomyces viridifaciens* ATTC11989 (accession CP023698) was used to determine the putative correct start and stop codons for *comEC*.

To create pIJ82-GFP, the region containing the *eGFP* gene with a *gap1* promoter was amplified from plasmid pGreen (Zacchetti et al., 2016) using primer pair gap1_FW_BglII and egfp_RV_EcoRI. The resulting PCR product was cloned into pIJ82 using BglII and EcoRI to generate the final plasmid.

### Construction of bacterial strains

To create new strains, transformation of L-form *alpha* with plasmid DNA was achieved using chemical transformation based on polyethylene glycol (PEG) (Kieser et al., 2000). Plasmid DNA was isolated from *Escherichia coli* ET12567/pUZ8002 to obtain methylation-deficient DNA. L-form strains *alpha* pIJ82-GFP and *alpha*Δ*divIVA* pIJ82-GFP were created using chemical transformation of *alpha* and *alpha*Δ*divIVA* with pIJ82-GFP respectively, followed by selection with hygromycin B (Table S2). The strains were verified using the detection of fluorescent eGFP production using fluorescence microscopy. Strain *alpha*Δ*comEA/EC* was obtained by chemical transformation of *alpha* with pRK1 followed by selection for apramycin (Table S2). Subsequent growth on non-selective medium allowed for double homologous recombination leading to replacement of the *comEA/EC* region by an apramycin resistance cassette, leading to thiostrepton-sensitive, apramycin-resistant cells. The strain was verified by PCR using primer pair ComEA_Apra_check_FW and ComEC_Apra_check_RV to confirm replacement of the region by the apramycin cassette. To further confirm deletion of this region, PCR was performed using primer pairs ComEC_Presence_Check_1_FW/RV and ComEC_Presence_Check_2_FW/RV, which amplify parts of *comEC* only if this genomic region is still present.

### Genomic DNA preparation

Genomic DNA was isolated from a 5-day old culture of *alpha*Δ*ssgB* (Ramijan et al., 2018) using phenol:chloroform extraction (Kieser et al., 2000). Briefly, the cell pellet was resuspended in 10.3% sucrose containing 0.01M ethylenediamine tetraacetic acid (EDTA) pH8 following lysis with 10% sodium dodecyl sulfate (SDS). Extraction with phenol:chloroform was performed and the nucleic acids were precipitated using isopropanol. The pellet was dissolved in Tris-EDTA buffer followed by RNase A (Thermo Fisher) and Proteinase K treatment (Qiagen). The gDNA was isolated using phenol:chloroform extraction and precipitated using absolute ethanol, before resuspension in nuclease-free water. Fragmented gDNA was obtained by beat-beating the intact gDNA for 12 minutes using 2 mm diameter glass beads in a Mikro-Dismembrator U (Sartorius) at 2000 rpm. Chromosomal DNA concentrations were verified using the Quant-IT™ Broad-Range dsDNA Assay Kit (Invitrogen).

### Preparation of protoplasts from *Kitasatospora*

*K. viridifaciens* strain DSM40239 was inoculated at a density of 5 x 10^6^ spores ml^-1^ in TSBS:YEME (1:1) liquid medium with 0.5% (w/v) glycine and 5 mM MgCl_2_. The culture was grown for 48 h while shaking at 200 rpm, after which protoplasts were prepared as described (Kieser et al., 2000). Cultures of 72 h were used for *K. viridifaciens* pIJ82-GFP and pRed*. Lysozyme treatment was performed by the addition of 10 mg ml^-1^ of chicken egg-white lysozyme (Sigma 70 000 U mg^-1^) to the mycelial suspension. The cells were incubated for 2-3 h at 100 rpm and 30°C, after which mycelial fragments were separated from the protoplasts by filtration trough a cotton wool filter (Kieser et al., 2000).

### Isolation of S-cells from *Kitasatospora*

S-cells were isolated from LPB cultures by filtration (Ramijan et al., 2018). In short, the culture was filtered through a sterile EcoCloth™ filter (Contec) and subsequently passed through a 5 µm Isopore™ membrane filter. The cells were concentrated by gentle centrifugation at 1000 xg for 20 minutes, after which 90% of the supernatant was removed. The cell pellet was suspended carefully in the remaining liquid.

### Chemical transformation

Polyethylene-glycol (PEG) was used for transformation as described (Kieser et al., 2000), using freshly prepared protoplasts, S-cells or L-forms that were kept on ice prior to transformation. For chemical transformation, 50 µl of cells were mixed with 1 µg pRed* (Zacchetti et al., 2018), 150 ng gDNA of strain *alpha*Δ*ssgB*, filter-sterilized salt-lysed cells (35 ng DNA from *alpha*Δ*ssgB*) or MilliQ. Then, 200 µl of 25% (w/v) PEG1000 in P-buffer (Kieser et al., 2000) was added to the cells, followed by gently mixing and diluting the suspension in P-buffer. Serial dilutions were plated on LPMA medium and after 16-18 h incubation an overlay was performed with 1 ml of P-buffer containing antibiotics. Colony forming units (CFU) were counted after 7 and 14 days for L-forms and S-cells/protoplasts, respectively. Transformants were verified by streaking on selective medium and microscopy.

### Natural transformation assay

Freshly prepared cells were incubated with 30 ng µl^-1^ DNA or MilliQ for 18-24 h at 100 rpm. A final concentration of 100 and 10 ng µl^-1^ intact gDNA and 10 ng ul^-1^ for fragmented gDNA isolated from *alpha*Δ*ssgB* was used in combination with both 1- and 7-day old *alpha*. Dilutions were plated on selective and nonselective LPMA after careful resuspension. Colony forming units were determined after 7-day incubation at 30°C for L-forms and mycelium and up to 14 days for protoplasts and S-cells. Transformants were verified by growth on selective medium and by PCR (using primers Tsr_Hyg_FW1 and Tsr_Hyg_RV1) or microscopy. Cells were prepared from at least five replica cultures to compare transformation efficiencies between strains.

### Membrane fluidity

Three replicate cultures of 1, 3 and 7-day old L-forms or freshly prepared protoplasts were subjected to a Laurdan dye assay as a measure for membrane fluidity (Scheinpflug et al., 2017). 1 ml of each culture was first centrifuged at 1000 xg for 10 minutes to remove any traces of the culture media. Cells were resuspended in 1 ml P-buffer and adjusted to an OD_600_ of 0.6. 10 mM Laurdan (6-Dodecanoyl-2-Dimethylaminonapthalene) stock solution (Invitrogen) was prepared in 100% dimethylformamide (DMF) and stored at -20°C in an amber tube. To each 1 ml OD-adjusted culture, 1 µl of Laurdan dye was added to a final concentration of 10 µM. The cultures were then incubated in the dark at 30°C for 10 min, while shaking at 100 rpm. The cells were washed three times with P-buffer containing 1% dimethyl sulfoxide to remove unbound dye molecules before the cells were resuspended in P-buffer. 200 µl of this resuspended culture was aliquoted into a 96-well black/clear glass bottom sensoplate (Greiner Bio-one VWR). Four technical replicas were measured per culture, as well as one replica per culture condition without dye to measure background fluorescence.

Sample excitation was performed at 350 nm followed by fluorescence emission capture at 435 and 490 nm, determined using a Spark® multimode microplate reader (Tecan). After subtracting the background fluorescence, the generalized polarization (GP) value was calculated using -

[math1]

Values obtained after calculation lie in the range of -1 to +1 with those closer to -1 indicating greater fluidity.

Preparation of cells for quantification of membrane fluidity by microscopy was performed as following. Cells were washed and OD-adjusted as mentioned above. Laurdan dye (stock concentration 10 mM) was added to 100 µl of culture to get a final concentration of 100 µM. The culture was placed in 30°C for 5 min, while shaking at 100 rpm in the dark. 900 µl of prewarmed P-buffer containing 1% dimethyl sulfoxide was added and the culture was centrifuged (1000 xg, 10 min) to remove any unbound dye molecules. The cells were finally resuspended in 100 µl of P-buffer for microscopy analysis. Cells treated similarly but without Laurdan dye were used a control for microscopy measurements.

### Preparation of fluorescently labelled DNA

Fluorescently labelled plasmid DNA was prepared using The Mirus Label IT^®^ Cy^™^5 Labelling Kit according to the manufacturer’s specifications. Aliquots of labelled DNA (100 ng µl^-1^) were stored at -20°C until further use.

### Self-assembly of lipid nanoparticles

All lipids (DLin-MC3-DMA/Cholesterol/DSPC/DMG-PEG2k/18:1 Liss Rhod PE) were combined in a molar ratio of 50/38.3/10/1.5/0.2 using stock solutions (100 µM – 10 mM) in chloroform:methanol (1:1). Organic solvents were evaporated under a nitrogen stream and remaining solvent was removed *in vacuo* for at least 1 h. Subsequently, the lipid film was dissolved in EtOH_abs_ and a 50 mM citrate buffer (pH = 4, MilliQ) was prepared. Each solution was loaded into separate syringes and connected to a T-junction microfluidic mixer. The solutions were mixed in a 3:1 flow ratio of citrate buffer against lipids (1.5 mL min^-1^ for citrate buffer, 0.5 mL min^-1^ for lipid solutions) giving a total lipid concentration of 1 mM. After mixing, the solution was directly loaded in a 10k MWCO dialysis cassette (Slide-A-Lyzer™, Thermo Scientific) and dialyzed against 1x Phosphate Buffered Saline (PBS, 137 mM NaCl, 2.7 mM KCl, 8 mM Na_2_HPO_4_ and 2 mM KH_2_PO_4_) overnight. All incubations with LNPs were performed with cells resuspended in LPB of which the final volume of LNP solution was 25%.

### Hydrodynamic diameter and zeta-potential measurement

Dynamic light scattering (DLS) measurements were performed on a Zetasizer Nano Series (Malvern Instruments, Malvern, UK). The incorporated HeNe laser works at a wavelength of 633 nm and uses a detector at an angle of 173° (noninvasive back scatter technology). Measurements were recorded with 1 min equilibration time in UV cuvettes at 25 °C. For the estimation of z-average diameter (intensity weight mean diameter) and polydispersity index (PDI)(relative width of particle size distribution) samples were prepared by tenfold dilution with 1x PBS. For the estimation of the zeta potential the sample was diluted with 0.1x Phosphate Buffered Saline (13.7 mM NaCl, 0.27 mM KCl, 0.8 mM Na_2_HPO_4_, and 0.2 mM KH_2_PO_4_). All the data were in triplicates to obtain the mean value.

### Fluorescence and light microscopy

Detection of fluorescence emission of transformants was performed using a Zeiss Axioscope A.1 equipped with a Zeiss Axiocam 305 color digital camera, using filter set 63 HE (Carl Zeiss, consisting of a 572/25 nm bandpass excitation filter, 590 nm beamsplitter and 629/62 nm bandpass emission filter) to capture mCherry fluorescence. All other microscopy was performed using a Zeiss LSM 900 confocal microscope with Airyscan 2 module, temperature control chamber and Zen 3.1 software (blue edition, Carl Zeiss Microscopy GmbH). All excitation and emission settings for this microscope are listed in Table S5. Multichannel (DIC and fluorescence) and multistack images were obtained unless specified otherwise. 10 μl of cells were imaged on an 8-chamber slide (ibidi®) coated with 0.1% poly-L-lysine (excess poly-L-lysine was removed and the slide was allowed to dry prior to applying the sample). For timelapse imaging or overnight incubation in the temperature control chamber, 400 μl of cell culture added to a 35 mm imaging μ-Dish (ibidi®) and allowed to settle at 30°C for an hour before overnight imaging. Image analysis was performed using Fiji (ImageJ) software (Schindelin et al., 2012).

Chromosomal DNA was visualized after incubation for 30 min with SYTO-9 at a final concentration of 2 μM. Cell membranes were visualized by incubation with SynapseRed C2M (SynapseRed) (PromoKine, PromoCell GmbH) at a final concentration of 0.2 μg ml^-1^. After overnight incubation in a μ-Dish (ibidi®) using the Zeiss LSM 900 confocal temperature control chamber, cells were imaged using the Airyscan mode with super resolution post-image processing via the Zen software. Protoplasts and S-cells were incubated with SynapseRed up to 72 h before imaging on a glass slide.

Uptake of fluorescently labelled DNA was assessed by incubating cells with Cy-5 labelled plasmid DNA (pFL-*ssgB*) at a final concentration of 1.25 μg ml^-1^ and was imaged after 48 h.

To capture internal vesicle formation and uptake of Dextran-Texas Red (D-TR), cells of *alpha* pKR2 were incubated with a final concentration of 1 mg ml^-1^ Dextran-Texas Red (3000 MW, neutral, Molecular Probes) in PBS and were imaged overnight. Multistack imaging across 6 μm total distance with 1.5 μm steps was done with an image captured every 10 minutes. Uptake of D-TR in *alpha*, protoplasts or S-cells was assessed after incubation up to 72 h.

Uptake of red fluorescent LNPs (LNP-LR) by *alpha* was visualized by imaging after overnight incubation in a μ-Dish (ibidi®) or after incubation for up to three days prior to imaging as indicated. Inhibition of LNP uptake was performed by incubation in the presence of 1-, 2.5- or 10-mM sodium azide (Sigma) or incubation at 4°C, and images were obtained using via the Zen software after 0, 24 and 48 h. To determine the subcellular localization of LNP-LR in *alpha* pIJ82-GFP, imaging was performed using the Airyscan mode with super resolution post-image processing and analyzed using the pixel intensity of the red (LNP-LR) and green (eGFP) channels using the Plot Profile tool in Fiji (ImageJ).

To measure the membrane fluidity, samples were excited using a 405 nm laser and images were captured at emissions of 430 nm and 500 nm. GP value was calculated using the ‘Calculate GP’ plug-in in Fiji (Vischer, 2016) to obtain a histogram of pixel counts over the range of -1 to +1. Briefly, the image is split into individual channels followed by background subtraction and setting the non-significant pixels to zero. The images are then assigned letters “A” and “B” to calculate A-B and A+B using the image calculator. Finally, a ratio of (A-B)/(A+B) is shown as an image where minimum pixel values are set to -1 (red) and maximum pixel values set to +1 (blue). Using the analyze histogram function a list of values is obtained and used for plotting the distributions of different samples.

### Cryo-correlative fluorescence and electron microscopy

#### High pressure freezing

7-day old L-form stain *alpha* pIJ82-GFP expressing cytoplasmic eGFP was adjusted to OD_600_ of 2 in fresh medium containing 25% (v/v) PBS and a final concentration of 17% sucrose. Cells were incubated for four days, during which cells settled to the bottom. A few microliters of the resuspended L-form pellet was sandwiched between HPF (High-Pressure-Freezing) carriers with 2 mm internal diameter (either 0.1 mm or 0.05 mm cavity, Art. 241 and Art. 390 respectively, Wohlwend) and tailor-made grid labeled, flat-sided finderTOP (Alu-platelet labelled, 0.3 mm, Art.1644 Wohlwend) to allow an imprint of a finder matrix on the amorphous ice (de Beer et al., 2021). The finderTOP was treated with 1% L-α-phosphatidylcholine (61755, Sigma) in ethanol (1.00983.1000, Supelco) before freezing. The samples were then high pressure frozen (Live µ, CryoCapCell) and stored in liquid nitrogen until imaging.

To improve correlation between cryo-light and cryo electron microscopy, the frozen samples were loaded into a universal cryo-holder (Art. 349559-8100-020, Zeiss cryo accessory kit) using the ZEISS Correlative Cryo Workflow solution, fit into the PrepDek® (PP3010Z, Quorum technologies, Laughton, UK). Here, the HPF carriers fits into a universal cryo-holder, which subsequently can be placed into an adaptor specific for cryo-light or cryo-electron microscopy.

#### Cryo-fluorescence imaging to detect regions of interests (ROI)

The frozen samples were imaged with a cryo-stage adaptor (CMS-196, Linkam scientific inc.) applied to an upright confocal microscope (LSM900, Zeiss microscopy GmbH) equipped with an Airyscan 2 detector. Overview images (Zeiss C Epiplan-Apochromat 5x/0.2 DIC) were made with reflection microscopy to visualize the gridded pattern on the ice surface. Next, medium-resolution Z-stack images (Zeiss C Epiplan-Apochromat 10x/0.4 DIC) were taken with a 488 nm laser (0.4%) with a voxel size of 0.15 µm x 0.15 µm x 1.18 µm. Using this resolution, cells of interest could be selected and Z-stack images were created (Zeiss C Epiplan-Neofluar 100x/0.75 DIC) using a 488 nm laser (4%), with a voxel size of 0.08 µm x 0.08 µm x 0.44 µm. In addition, the ice surface was imaged in all ROIs with reflection microscopy for correlation purposes in the FIB-SEM.

Prior to cryo-light imaging, a Zeiss ZEN Connect project (Zeiss software for correlative microscopy, version 3.1) was created to make a working sheet (canvas) to align and overlay all the images and to facilitate further correlation with cryo-FIB-SEM.

#### 3D Cryo-FIB-SEM

The sample was sputter-coated with platinum, 5mA current for 30 seconds, using the prep stage sputter coater (PP3010, Quorum technologies, Laughton, England) and was transferred into the Zeiss Crossbeam 550 FIB-SEM (Carl Zeiss Microscopy GmbH, Oberkochen, Germany) using the PP3010T preparation chamber (Quorum, Laughton, England). Throughout imaging, the samples were kept at -140 °C and the system vacuum pressure was 1 x 10^−6^ mbar.

After inserting the sample into the FIB-SEM chamber, overview images were taken using the SEM to align the data with the LSM reflection image of the surface of the same ZEN Connect project. This alignment enables the stage registration which allows using the fluorescence signal to navigate to different regions of interest. After initial alignment using the SEM, a FIB image of the surface was collected with the 30kV@10pA probe in 54° tilt.

A coarse trench was milled for SEM observation using the 30 kV@30 nA FIB probe. Cold deposition was done with platinum for 30 sec. Fine FIB milling on the cross section was done using the 30kV@700pA probe. For serial FIB milling and SEM imaging the slice (trench) width was 40 μm and for FIB milling the 30 kV@300pA probe was used, with a slice thickness of 20 nm. When a new slice surface was exposed by FIB milling, an InLens secondary and EsB images were simultaneously collected at 2.33 kV acceleration potential with 250pA probe current. The EsB grid was set to -928 V. The image size was set to 2048 × 1536 pixels. For noise reduction line average with a line average count N = 46 at scan speed 1 was used. The voxel size of all stacks was 5×5×20 nm^3^.

#### 3D FIB-SEM Image post processing

The cryo-FIB-SEM images were processed using MATLAB (R2018b, Natick, Massachusetts: The MathWorks Inc.) to correct for defects such as curtaining, misalignment and local charging. The same software was used for subsequent noise reduction and contrast enhancement. A summary of each processing step is as follows:

***Curtaining:*** Removing the vertical stripes in the stacks was done following a wavelet-FFT filtering approach described by (Munch et al., 2009). In brief, the high frequency information corresponding to the vertical stripes was successively condensed into a single coefficient map using decomposition by “coif” wavelet family. Subsequently, a 2D-fourier transform was performed to further tighten the stripe information into narrow bands. Finally, the condensed stipe information was eliminated by multiplication with a gaussian damping function and the destriped image was reconstructed by inverse wavelet transform.

***Alignment:*** The consecutive slices were aligned using normalized cross correlation. Briefly, the first image in the stack was chosen as reference and the second image was translated pixel by pixel across the reference and a normalized cross correlation matrix was obtained using the “normxcorr2” function. The location of the highest peak in the cross-correlation matrix (representing the best correlation) was then used to calculate the translation required to align the two images. Once the moving image was aligned with the reference image, it served as the reference for alignment of the subsequent slice.

***Charging:*** Elimination of the local charge imbalance was achieved using anisotropic gaussian background subtraction. Briefly, the “imgaussfilt” function was used to perform 2D-gaussian smoothing with a two-element standard deviation vector. The elements in the vector were chosen in a manner to apply a broad and sharp gaussian in the horizontal and vertical directions, respectively. Subsequently, the corrected image was obtained by subtracting the filtered image from the original image.

***Noise Reduction:*** In order to improve the signal-to-noise ratio, noise reduction was performed using anisotropic diffusion filtering (Perona and Malik, 1990). Briefly, using the “imdiffuseest” function, the optimal gradient threshold and number of iterations required to filter each image was estimated. Subsequently, the “imdiffusefilt” function was applied with the estimated optimal parameter values to denoise each image.

Contrast enhancement: As the final processing step, the contrast was enhanced using “Contrast-limited adaptive histogram equalization” (Zuiderveld, 1994). Using the “adapthisteq” function, the contrast was enhanced in two steps, using a uniform distribution and a low clipping limit in order to avoid over-amplification of homogeneous regions.

***3D segmentation:*** DragonflyTM image analysis and deep-learning software (version 2021.1, Objects Research Systems, Montreal, QC, Canada) was used to segment all image data.

### Bioinformatic search for putative competence genes

Protein sequences from *Bacillus subtilis* str. 168, *Neisseria gonorrhoeae* and *Helicobacter pylori* strain P12 were obtained from the UniProt database or literature (Wolfgang et al., 1999). These sequences were used for a BlastP search against the non-redundant protein sequence database of *Streptomyces viridifaciens* (taxid 48665). Hits belonging to *Streptomyces viridifaciens* strain DSM40239, sequence accession numbers CP090840 to CP090842 with an E-value of 1×10^-6^ or lower were collected (Table S1).

### Statistics

All statistics were performed using SPSS statistics software (IBM, version 27.0). P-values less than 0.05 were considered statistically significant.

## SUPPLEMENTAL VIDEO LEGENDS

**Video S1. Uptake of Dextran-Texas Red by L-forms, Related to Figure 2**

Timelapse video of *alpha*-DivIVA-eGFP (green) incubated with 3 kDa Dextran-Texas Red (D-TR; magenta). Left: Brightfield. Right: Composite of green and magenta channels. Scale bar indicates 1 µm.

**Video S2. 3D Reconstruction of Vesicles in L-form Cell, Related to Figure 4**

3D segmentation of *alpha* pIJ82-GFP corresponding to Figure 4 Jiv and Kiv. Colours indicate individual vesicles or vesicle complexes. The cell is depicted in grey.

**Video S3. 3D Reconstruction of Vesicles in L-form Cell, Related to Figure 4**

3D segmentation of *alpha* pIJ82-GFP corresponding to Figure 4H and Figure S6D. Colours indicate individual vesicles or vesicle complexes. The cell is depicted in grey.

## ACKNOWLEDGEMENTS

R.K. and L.Z. are supported by the TARGETBIO program of the Netherlands Organization for Scientific Research (NWO), grant nr. 15812. S.S. is supported by the NWA startimpulse grant (Origins Centre). As part of the COFUND project oLife, D.A. acknowledges funding from the European Union’s Horizon 2020 research and innovation program under the Grant Agreement 847675. M.d.B, R.R., D.D. and A.A. are supported by an ERCAdvanced Investigator grant (H2020-ERC-2017-ADV-788982-COLMIN). A.A. is also supported by the NWO (VI.Veni.192.094). We thank Nico Sommerdijk (Radboudumc, Electron Microscopy Center) for his contribution with high pressure freezing of the samples.

---

**Figure S1.**
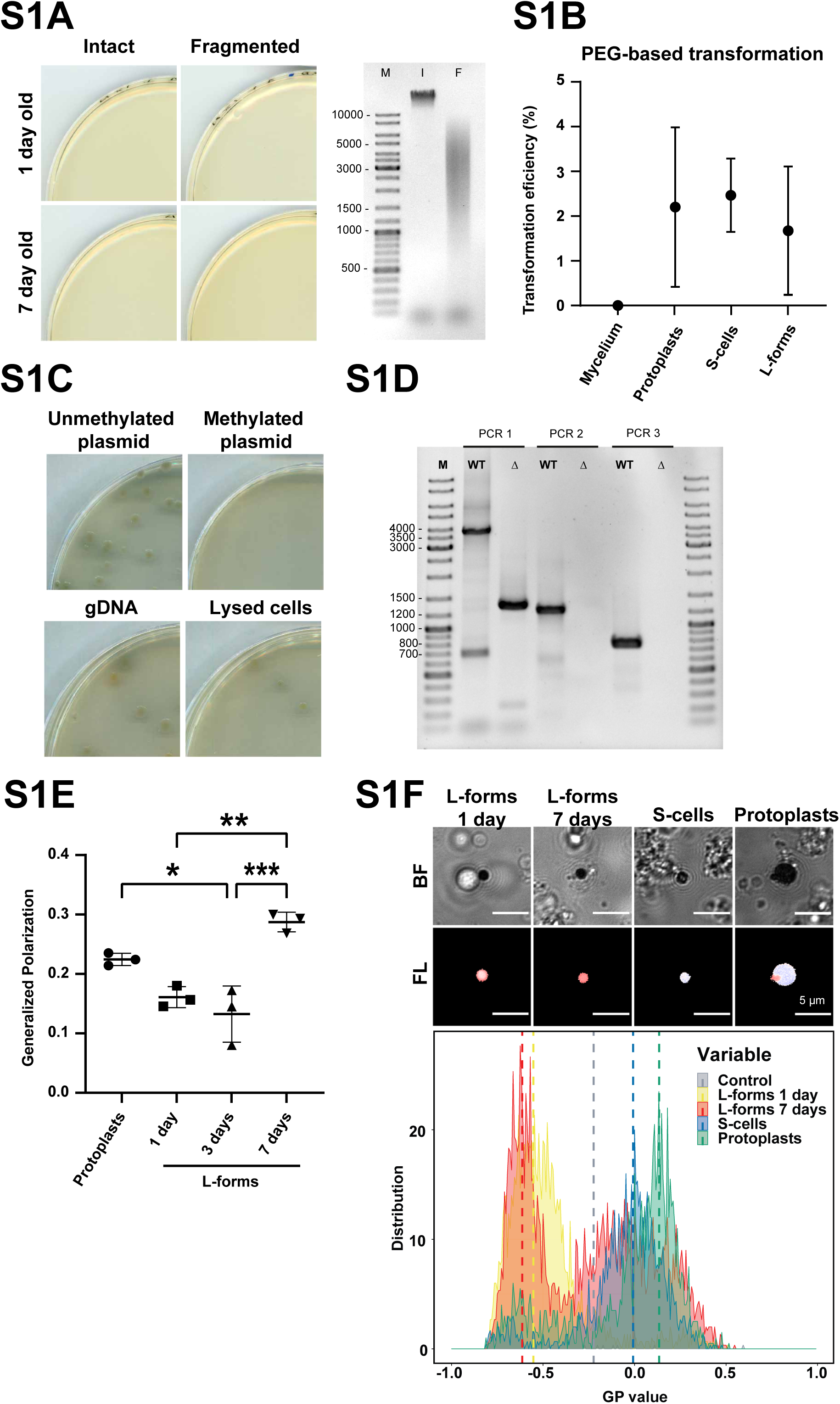
Analysis of Natural and Artificial DNA Uptake and Membrane Fluidity of Cell-Wall Deficient Cells and Confirmation of alphaΔcomEA/EC Mutant, Related to Figure 1. (A) (Left) Transformation plates showing absence of natural transformation upon incubation of 1-and 7-day old L-form alpha with intact or fragmented gDNA of *alphaΔssgB* containing an apramycin resistance cassette. (Right) Gel electrograph of 100 ng intact (I) or fragmented (F) gDNA of *alphaΔssgB* as used in the natural transformation assay. (B) Polyethylene glycol (PEG)-based transformation efficiency of *K. viridifaciens* mycelium, protoplasts, S-cells and L-forms using plasmid DNA (pRed*) containing an apramycin resistance gene, shown as the percentage of trans-formed colonies per total colony forming units. Data are represented as mean ± SD, n=3. (C) PEG-based transformation of alpha using unmethylated or methylated plasmid DNA (pRed*), gDNA or filter-sterilized salt-lysed cells from mutant line *alphaΔssgB*. (D) Gel electrograph of PCR products from three different PCR mixes to confirm the replacement of *comEA* and *comEC* by an apramycin resistance cassette. WT = gDNA *alpha*; Δ = gDNA *alphaΔcomEA/EC*. Expected products: PCR 1 WT = 3676 bp, mutant = 1294 bp; PCR 2 WT = 1197 bp, mutant = no amplification, PCR 3 WT = 745 bp, mutant = no amplification. (E) Generalized Polarization (GP) as measure of membrane fluidity of *K. viridifaciens* protoplasts, 1-, 3- and 7-day old L-form *alpha*. Lower GP indicates higher fluidity. *, ** and *** indicate P ≤ 0.05, 0.01 and 0.001, respectively (one-way ANOVA, F (3,8) = 19.49, Tukey post-hoc test, n=3). Data are represented as mean ±SD with individual data points, n=3. (F) Membrane fluidity of L-form *alpha* (1- and 7-day old), S-cells and protoplasts of *K. viridifaciens*. Top rows show brightfield images and heatmap of fluorescence emission (red to blue colour indicate GP values of -1.0 to 1.0 respectively) of representative cells stained with a Laurdan dye for quantifying the membrane fluidity (BF = brightfield, FL = fluorescence emission). Bottom panel shows frequency distributions of the Generalized Polarization (GP). Lower GP values correspond to higher membrane fluidity indicating that L-forms have more fluid membranes compared to S-cells and protoplasts. Control = cells imaged and analysed without Laurdan staining.

**Figure S2.**
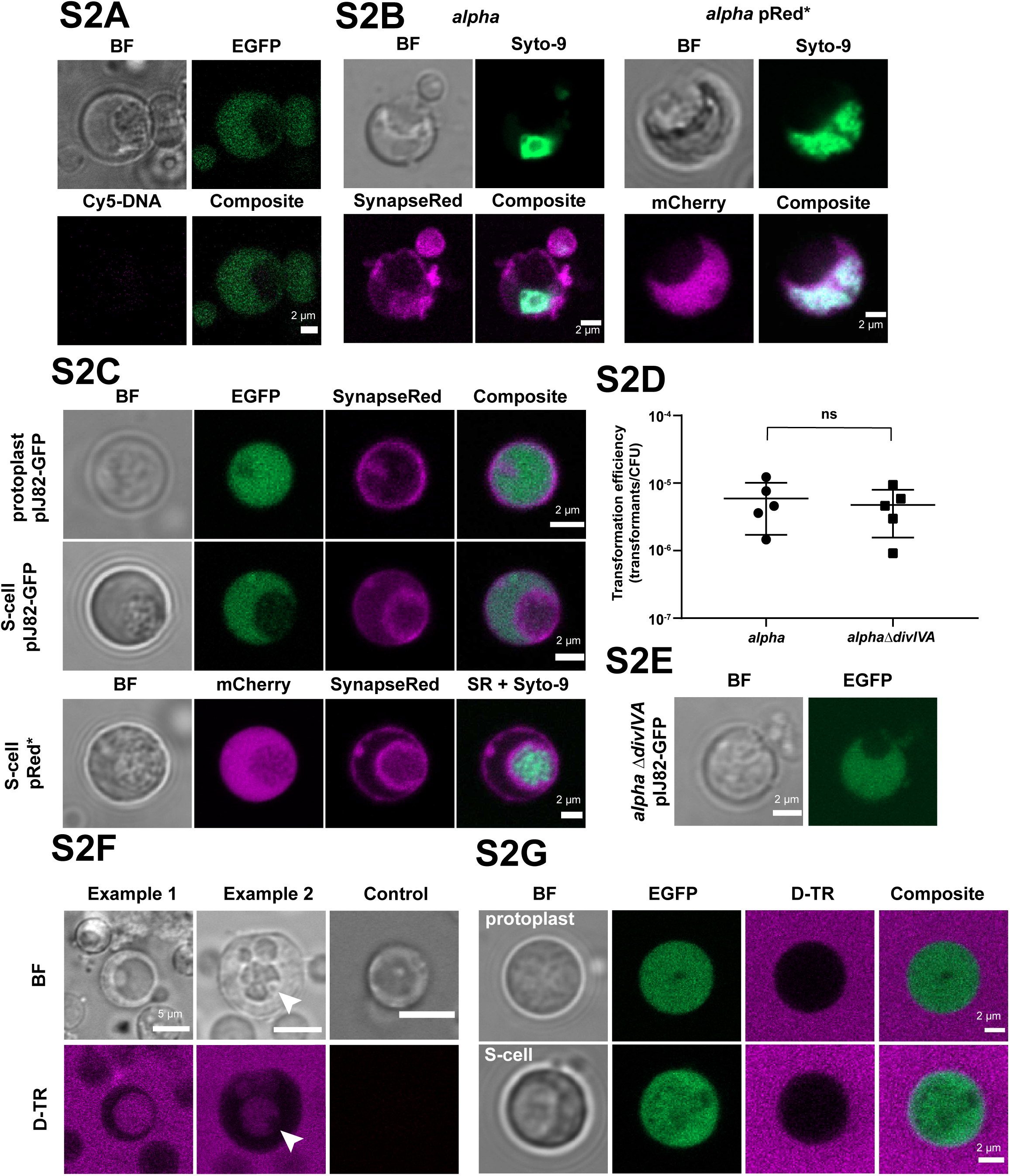
Analysis of DNA Content, Internal Vesicles and Uptake of D-TR of Cell-Wall Deficient Cells, and effect of divIVA deletion on DNA Uptake, Related to Figure 2. (A) *alpha* pIJ82-GFP incubated without Cy-5 DNA as fluorescence control. (B) *alpha* and *alpha* pRed* stained with SYTO-9 (green) to indicate chromosomal DNA. *alpha* is stained with SynapseRed C2M (SynapseRed; magenta) to visualize cell membranes, whereas (absence of) cytosolic mCherry for *alpha* pRed* (magenta) indicates the presence of an internal vesicle. (C) Protoplasts and S-cells of *K. viridifaciens* pIJ82-GFP producing cytosolic eGFP incubated with SynapseRed for 72 h (top rows), and S-cells of *K. viridifaciens* pRed* producing cytosolic mCherry incubated with SynapseRed (SR) and SYTO-9 for 72 h (bottom row). Chromosomal DNA is visualized using SYTO-9 staining. Note that presence of internal membrane structures causes a reduction in cytosolic fluorescence emission. (D) Natural transformation assay of 7-day old *alpha* and *alphaΔdivIVA* using pFL-*ssgB*. ns = not significant (two-tailed independent t-test, t(8)=0.489, P=0.638). Data are represented as mean ± SD with individual data points, n=5. (E) L-forms without DivIVA can produce internal vesicles as shown for 5-day old *alphaΔdivIVA* pIJ82-GFP producing cytosolic eGFP. Scale bar = 2 μm. (F) *alpha* incubated with (example 1 and 2) or without (control) Dextran Texas-Red (D-TR; magenta), showing the formation of internal vesicles filled with D-TR. The arrow indicates the presence of a non-fluorescent secondary internal vesicle inside an existing internal vesicle (example 2). Scale bar = 5 μm. (G) Protoplasts and S-cells of *K. viridifaciens* pIJ82-GFP incubated with D-TR for 72 h. Note that no internalization of D-TR was observed.

**Figure S3.**
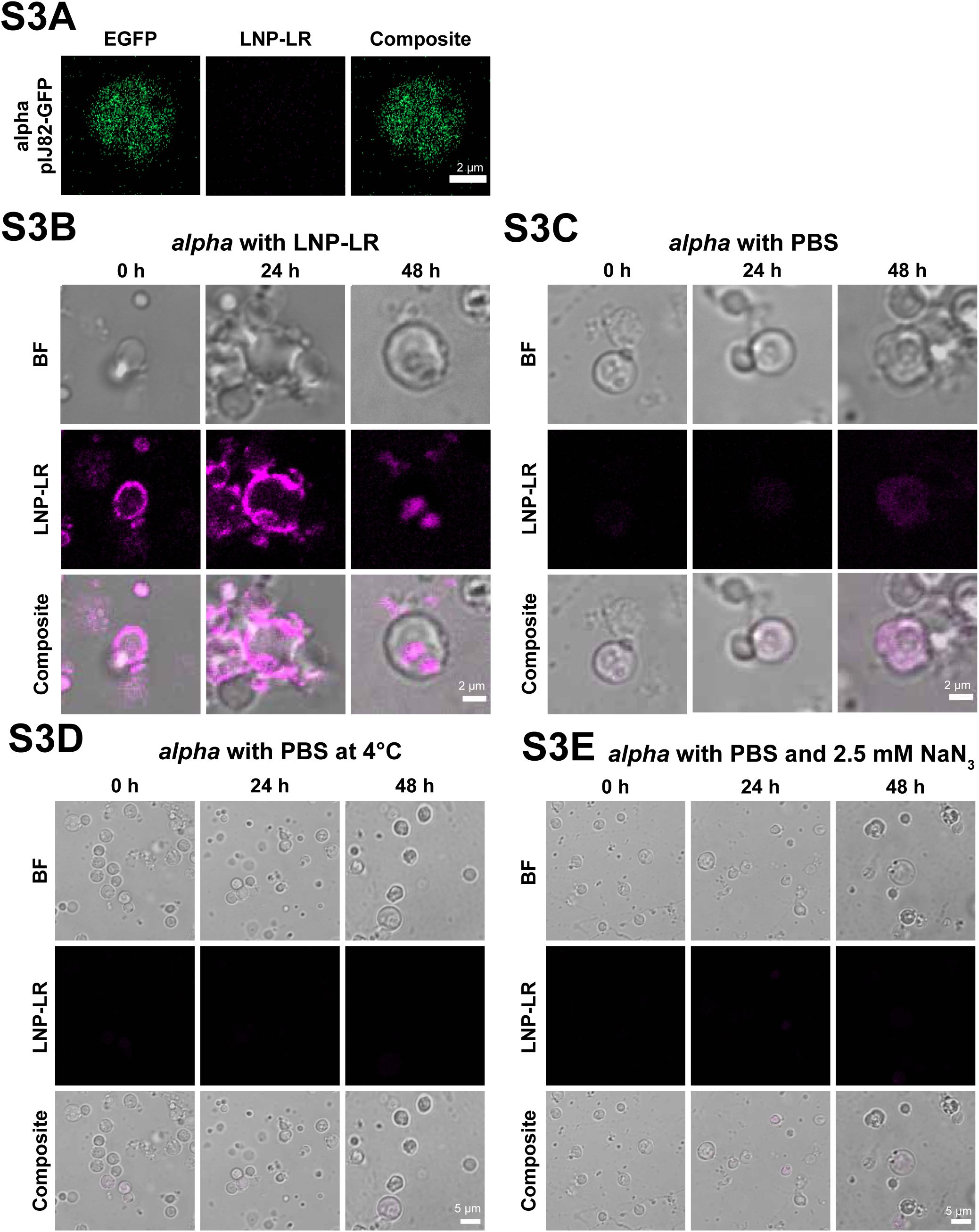
Uptake of LNP-LR by alpha, Related to Figure 3. (A) *alpha* pIJ82-GFP incubated without LNP-LR (LNP-Liss Rhod; magenta) as imaging control. (B-C) *alpha* incubated with (B) or without (C) LNP-LR showing localization of LNP-LR after 0 h, 24 h and 48 h or examples of autofluorescence, respectively. (D-E) *alpha* incubated with PBS at 4 degrees (D) or with PBS at 30°C in the presence of 2.5 mM sodium azide (E) as control for fluorescence emission. Images were obtained after 0, 24 and 48 h incubation.

**Figure S4.**
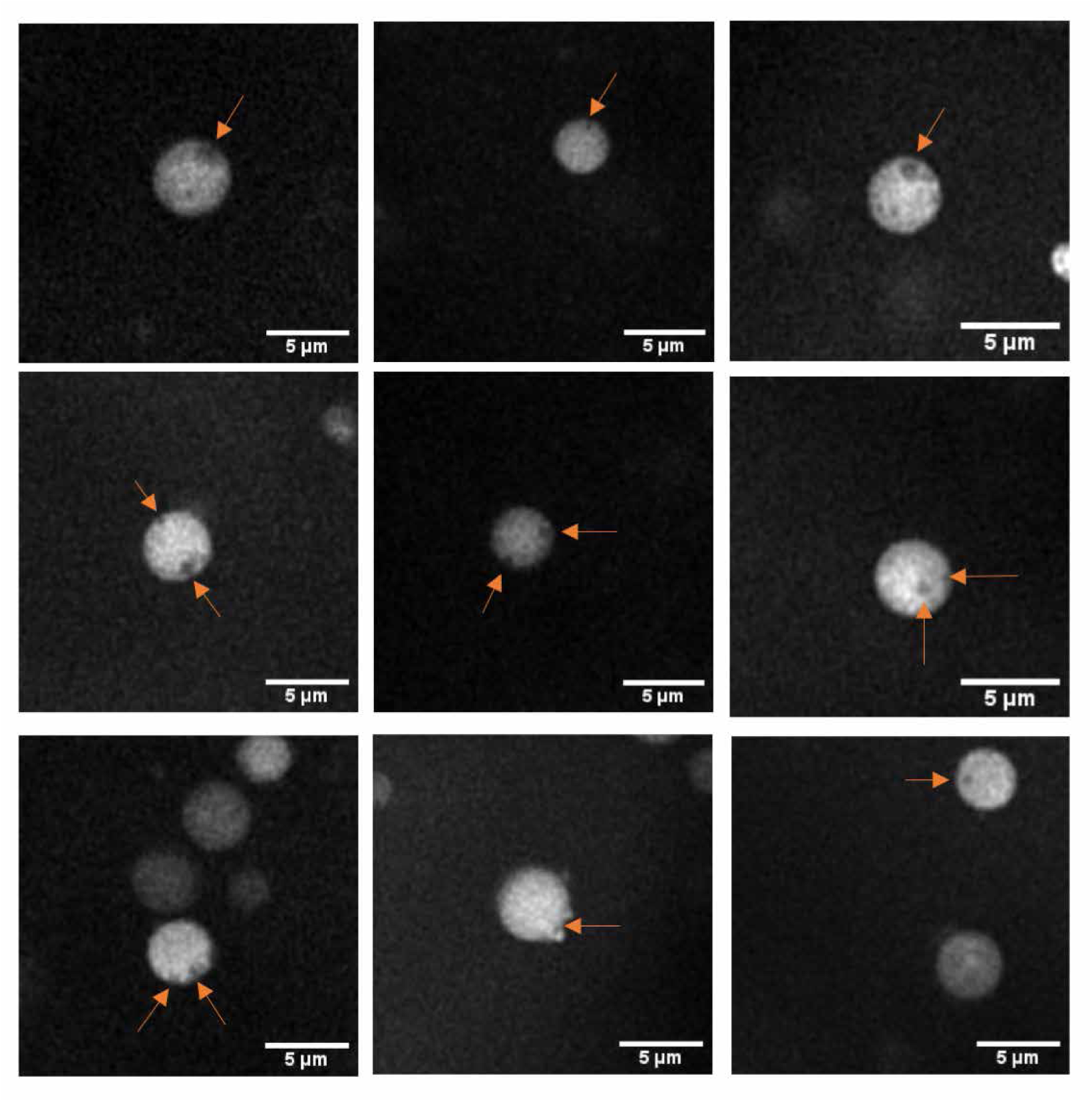
High Resolution Cryo-Fluorescence of L-forms, Related to Figure 4. *alpha* pIJ82-GFP imaged using cryo-fluorescence microscopy. Putative vesicles are indicated with arrows. Images were captured using the long distance 100x objective.

**Figure S5.**
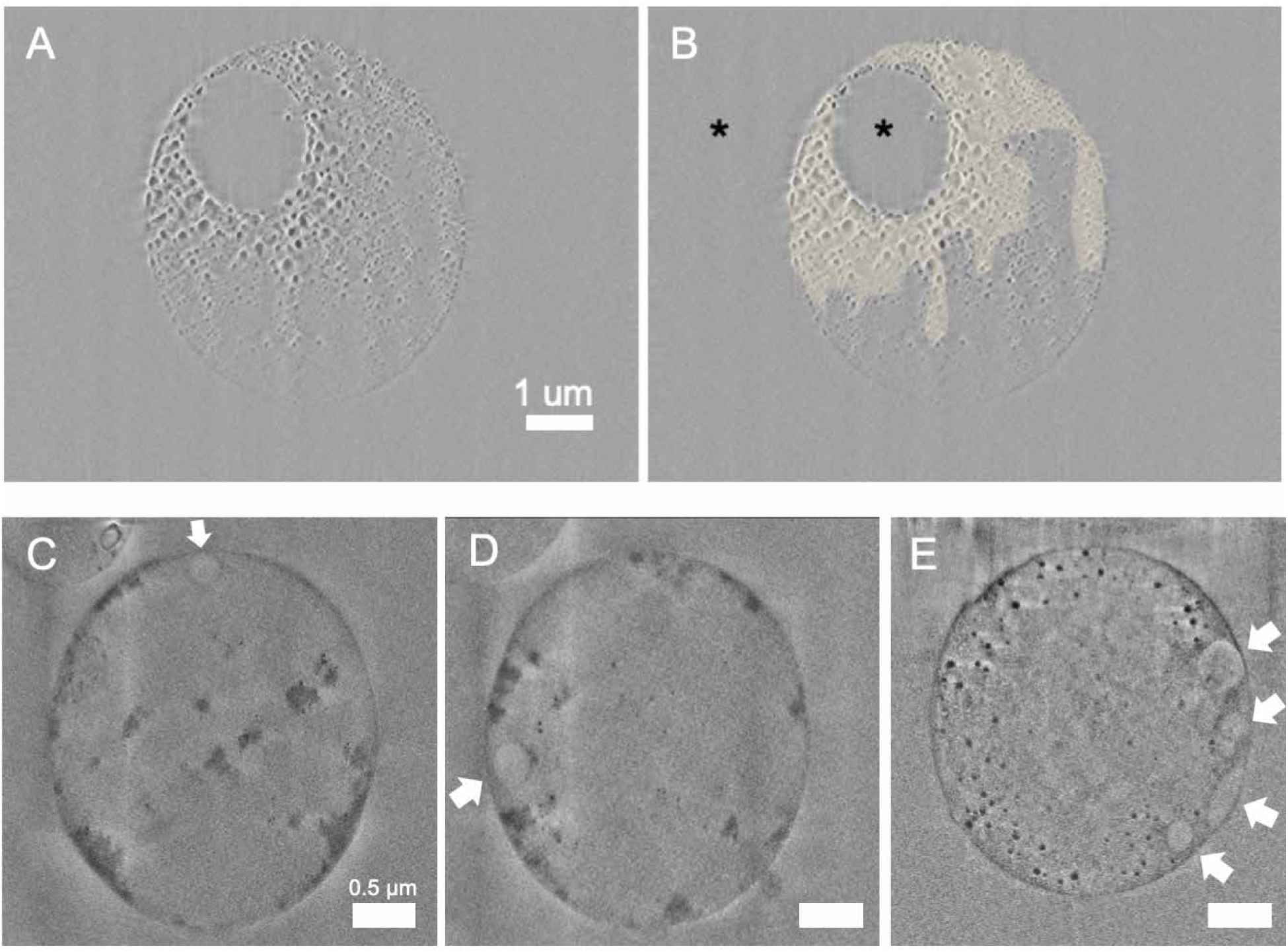
Over-dose experiment of L-form Cell using FIB-SEM, Related to Figure 4. (A-B) FIB-SEM slice of over-dose experiment using *alpha* pIJ82-GFP. The yellow colour in B) indicates areas with distinguished beam damage. The vesicle (black asterisk in the center of the cell) seems to be less to none affected by the dose, similar to the medium outside the cell (black asterisk outside of the cell). The image in Figure 4D is taken before this experiment, and Figure 4E is obtained by summing several slices deeper in the cell after acquiring this image. (C-E) FIB-SEM slices of two cells (C-D correspond to the cell in Figure 4F and E corresponds to the cell in Figure 4 H-K), white arrows indicate vesicles that line the cell membrane. Scale bar in C-E is 0.5 μm.

**Figure S6.**
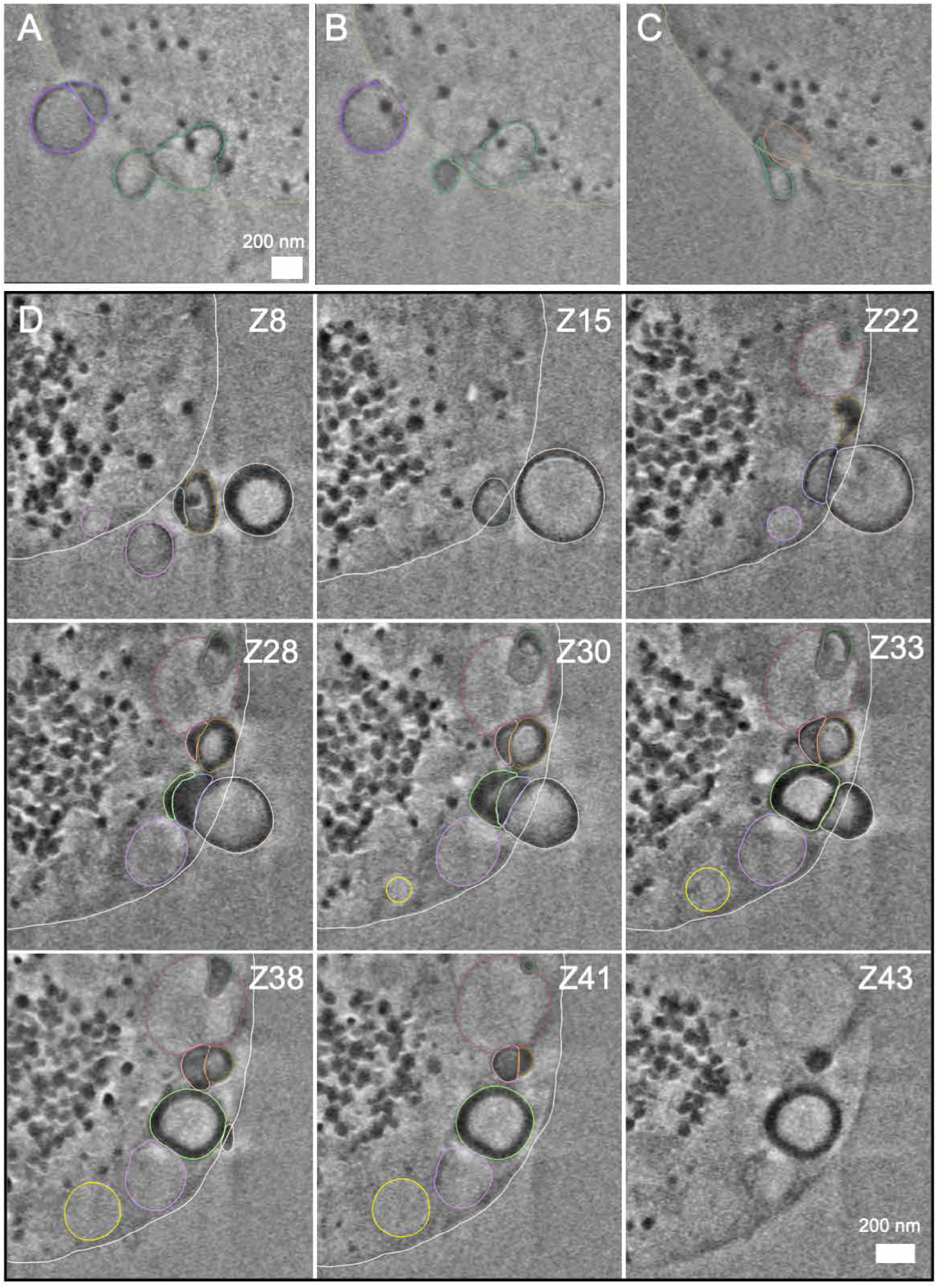
3D Segmentation of L-form Vesicles, Related to Figure 4. (A-C) FIB-SEM slices corresponding to Figure 4I, Jiv and Ki-iii, respectively. Colours correspond to the segmented colours in Figure 4Kiv. Vesicles that are budding out the cells are connected to other vesicles or are elongated inside the cell. Scale bar = 200 nm. See also Video S2. (D) FIB-SEM slices corresponding to the cell in Figure 4H. Z-number indicates the slice. Colours indicate individual vesicles. See also Video S3.

**Table S1.**
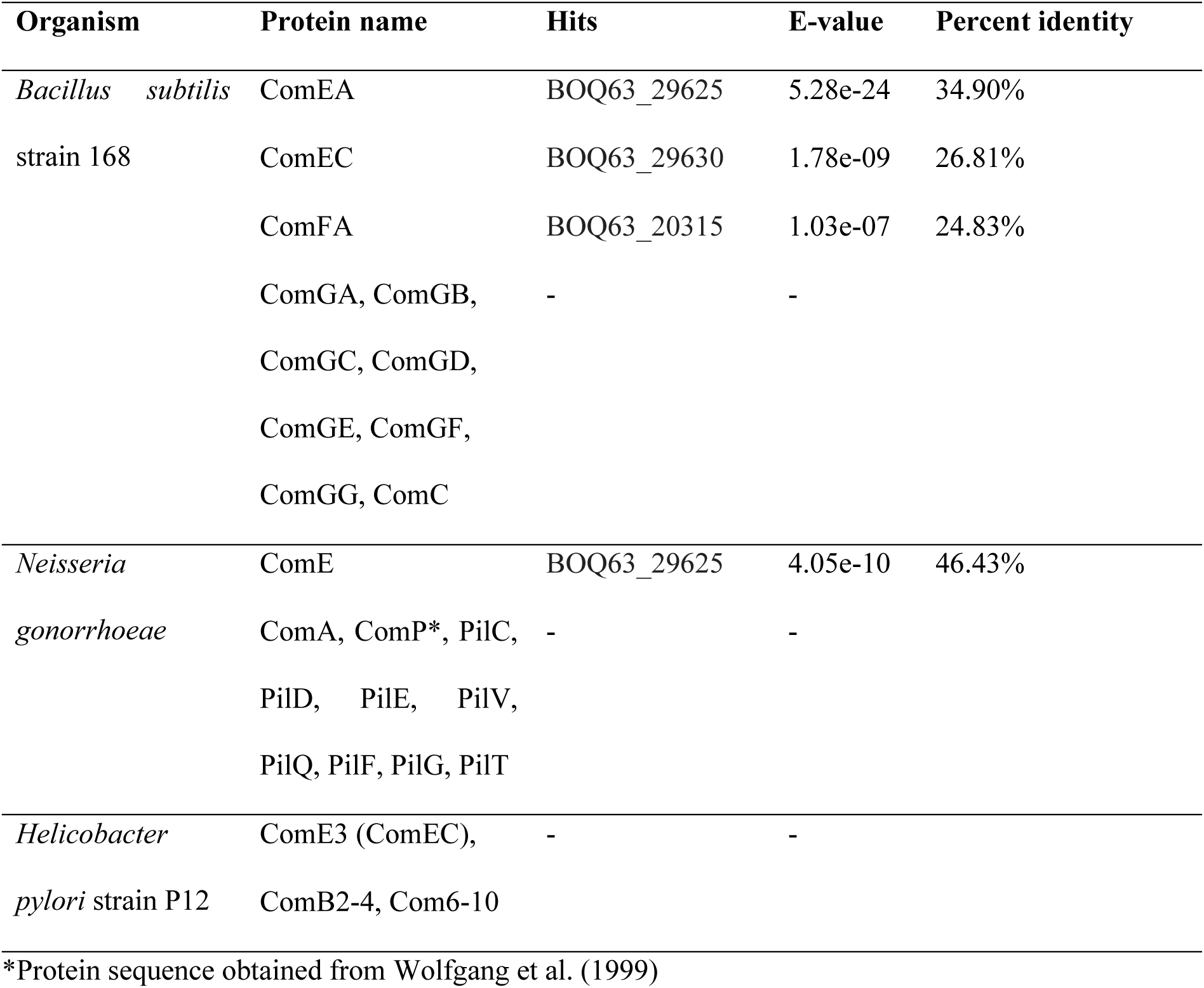
BlastP results showing significant hits (E-value < 1e-06), E-value and percent identity against *K. viridifaciens* DSM40239.

**Table S2.**
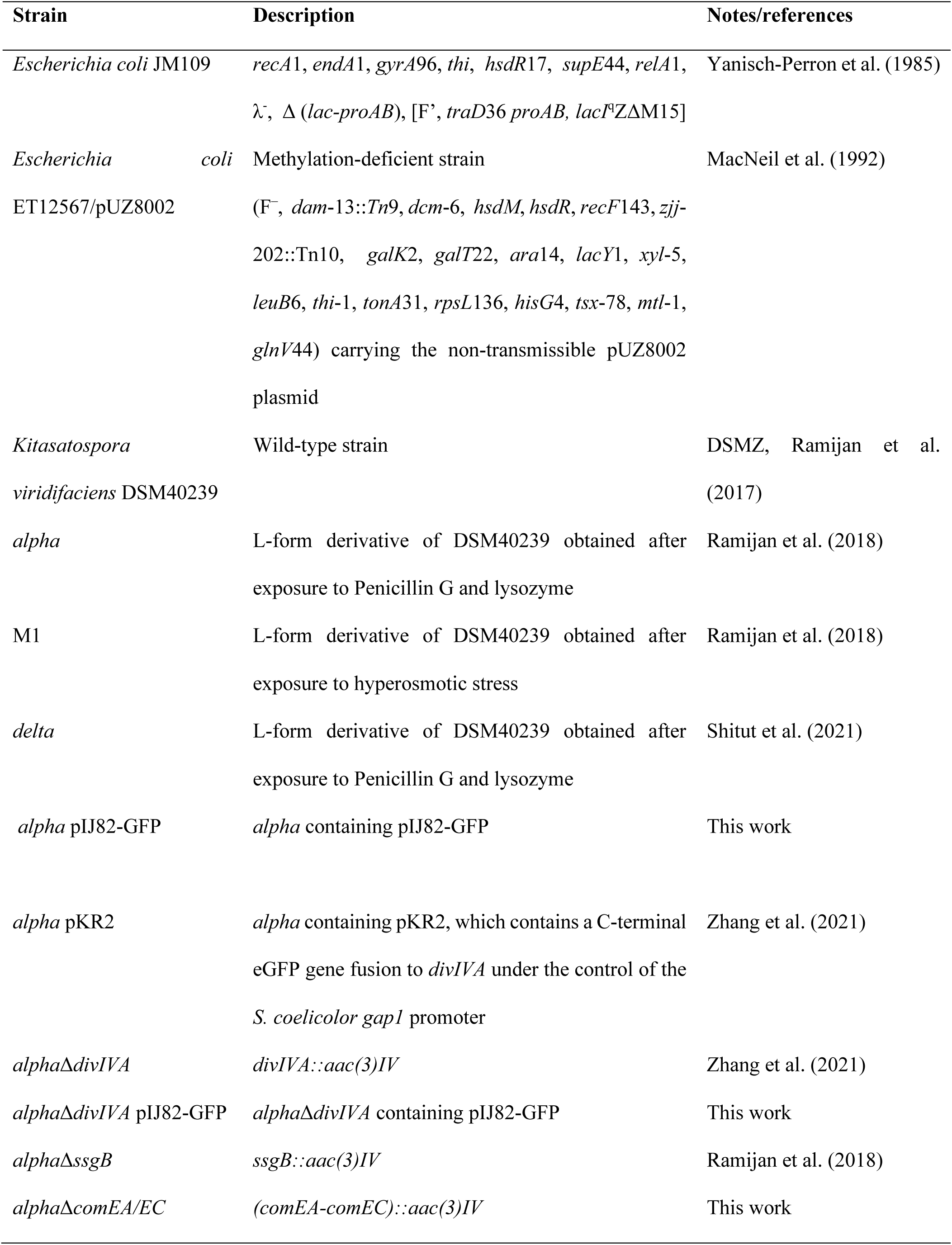
Strains used in this study.

**Table S3.** Plasmids used in this study.

| Plasmid | Features | Notes/References |
| --- | --- | --- |
| pRed* | pIJ8630-derivative expressing *mCherry* under control of the *S. coelicolor* A3(2) *gapI* promoter | Zacchetti et al. (2018) |
| pGreen | pIJ8630-derivative expressing *eGFP* under control of the *S. coelicolor* A3(2) *gapI* promoter | Zacchetti et al. (2016) |
| pSET152 | *E. coli-Streptomyces* shuttle vector; high copy number in *E. coli* and integrating in the ϕC31 *attB* site in *Streptomyces* | Bierman et al. (1992) |
| pIJ82 | pSET152-derivative carrying a hygromycin resistance cassette | Kindly provided by Dr. B. Gust (JIC) |
| pIJ82-GFP | pSET152-derivative expressing *eGFP* under control of the *S. coelicolor* *gapI* promoter | This work |
| pMS82 | *E. coli-Streptomyces* shuttle vector integrative in the ϕBT1 *attB* site for genomic integration in *Streptomyces* | Gregory et al. (2003) |
| pWHM3-oriT | Self-replicating, multi-copy, unstable plasmid harboring *oriT*, used as *E. coli/Streptomyces* shuttle vector | Wu et al. (2019) |
| pWHM3-oriT-hyg | pWHM3-oriT-derivative carrying a hygromycin resistance cassette inserted in to the *tsr* gene in the EcoRV site | This work |
| pFL-*ssgB* | pWHM3-oriT-hyg-derivative containing a hygromycin resistance gene and a downstream flanking sequence of *ssgB* downstream derived from pKR1 to enable integration into the genome | This work |
| pRK1 | pWHM3-oriT containing both flanks of the *comEA-comEC* region interspersed with the *apra-loxP* cassette conferring resistance to apramycin | This work |
| pKR1 | pWHM3-based construct used to replace *ssgB* by *aac(3)IV* | Ramijan et al. (2018) |
| pKR2 | pIJ8630 derivative carrying a viomycin resistance cassette and expressing a C-terminal *eGFP* fusion to *divIVA* under control of the *S. coelicolor* A3(2) *gapI* promoter | Zhang et al. (2021) |

**Table S4.**
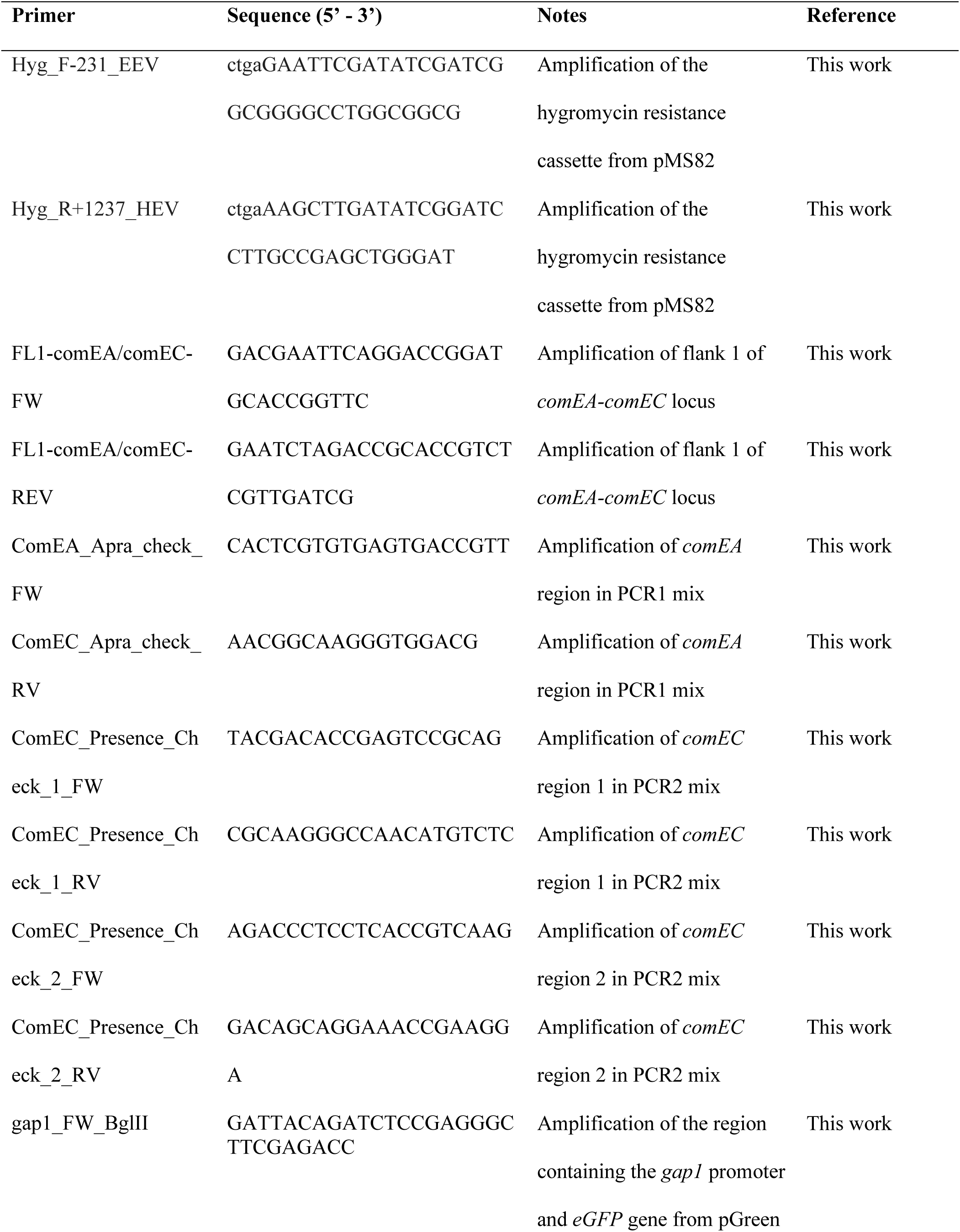
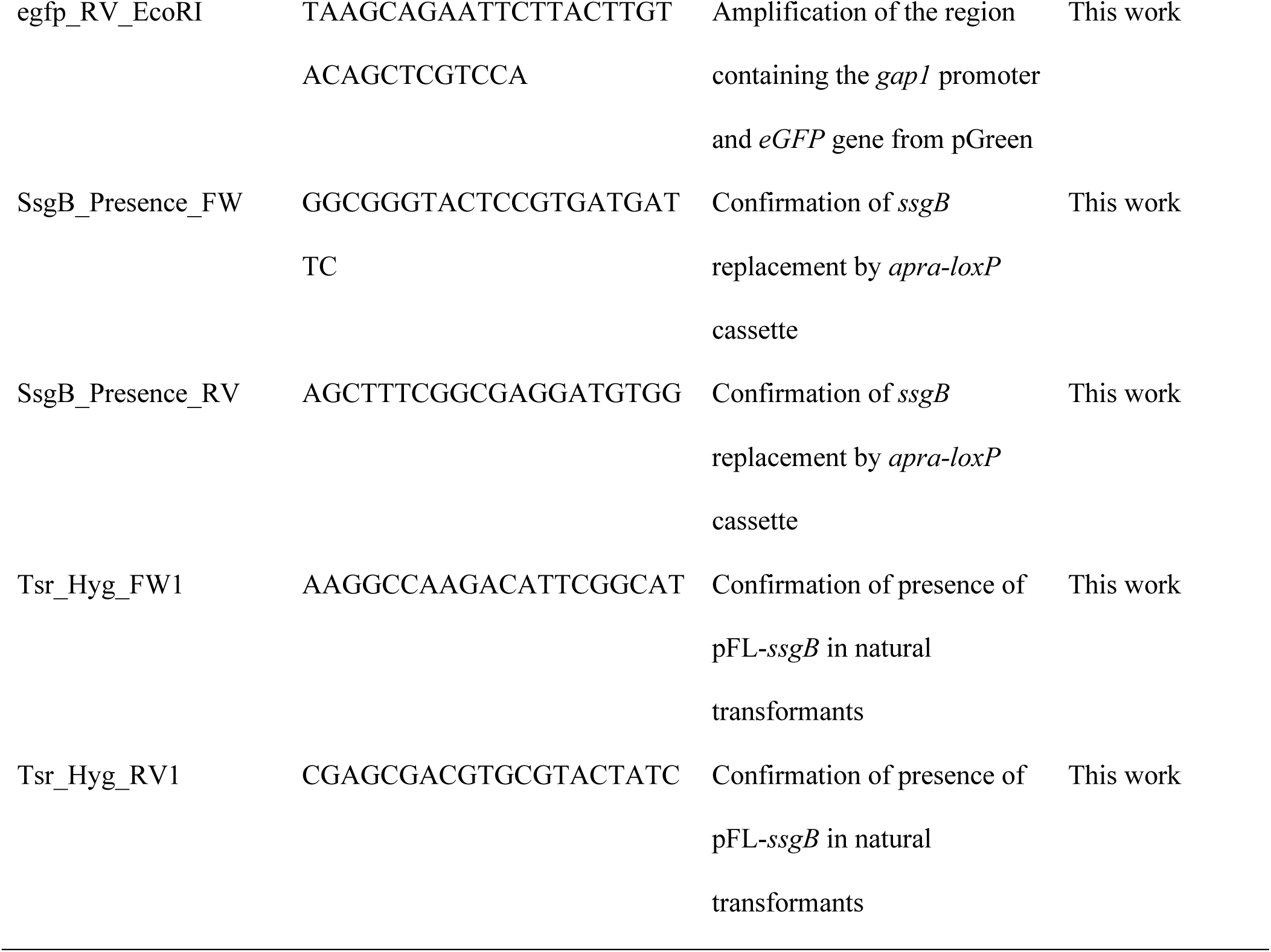
Primers used in this study.

**Table S5.** Imaging settings used with the Zeiss LSM 900 confocal microscope.

| Fluorescent dye or particle | Excitation (nm) | Emission filter (nm) |
| --- | --- | --- |
| eGFP | 488 | 490-575 |
| mCherry | 561 | 565-700 |
| SYTO-9 | 488 | 490-575 |
| SynapseRed C2M | 506 | 571-700 |
| Dextran-Texas Red | 584 | 560-700 |
| Cy5 | 650 | 450-700 |
| LNP-LR | 587 | 565-700 |

**Table S6.** Dynamic Light Scattering (DLS) and ζ-potential of lipid nanoparticles. PDI = polydispersity index.

| LNP | Avg. size (nm) at<br>25 °C | PDI | ζ-potential at 25<br>°C |
| --- | --- | --- | --- |
| LNP-LR | 125.8 | 0.126 | -8.5 |

## REFERENCES

1. Allan, E.J., Hoischen, C., and Gumpert, J. (2009). Bacterial L-forms. Adv Appl Microbiol 68, 1–39.

2. Araki, N., Johnson, M.T., and Swanson, J.A. (1996). A role for phosphoinositide 3-kinase in the completion of macropinocytosis and phagocytosis by macrophages. Journal of Cell Biology 135, 1249–1260.

3. Arnold, B.J., Huang, I.T., and Hanage, W.P. (2021). Horizontal gene transfer and adaptive evolution in bacteria. Nat Rev Microbiol.

4. Atkinson, H.A., Daniels, A., and Read, N.D. (2002). Live-cell imaging of endocytosis during conidial germination in the rice blast fungus, Magnaporthe grisea. Fungal Genetics and Biology 37, 233–244.

5. Behr, J.-P. (1997). The proton sponge: a trick to enter cells the viruses did not exploit. CHIMIA International Journal for Chemistry 51, 34–36.

6. Bendezu, F.O., and de Boer, P.A. (2008). Conditional lethality, division defects, membrane involution, and endocytosis in mre and mrd shape mutants of Escherichia coli. J Bacteriol 190, 1792–1811.

7. Bierman, M., Logan, R., O’Brien, K., Seno, E.T., Rao, R.N., and Schoner, B.E. (1992). Plasmid cloning vectors for the conjugal transfer of DNA from *Escherichia coli* to *Streptomyces* spp. Gene 116, 43–49.

8. Briers, Y., Staubli, T., Schmid, M.C., Wagner, M., Schuppler, M., and Loessner, M.J. (2012a). Intracellular vesicles as reproduction elements in cell wall-deficient L-form bacteria. PLoS One 7, e38514.

9. Briers, Y., Walde, P., Schuppler, M., and Loessner, M.J. (2012b). How did bacterial ancestors reproduce? Lessons from L-form cells and giant lipid vesicles: multiplication similarities between lipid vesicles and L-form bacteria. Bioessays 34, 1078–1084.

10. Budker, V., Budker, T., Zhang, G., Subbotin, V., Loomis, A., and Wolff, J.A. (2000). Hypothesis: naked plasmid DNA is taken up by cells in vivo by a receptor-mediated process. The journal of gene medicine 2, 76–88.

11. Cervia, L.D., Chang, C.C., Wang, L., and Yuan, F. (2017). Distinct effects of endosomal escape and inhibition of endosomal trafficking on gene delivery via electrotransfection. PLoS One 12, e0171699.

12. Chapman, D. (1975). Phase transitions and fluidity characteristics of lipids and cell membranes. Quarterly reviews of biophysics 8, 185–235.

13. Chen, I., and Dubnau, D. (2004). DNA uptake during bacterial transformation. Nat Rev Microbiol 2, 241–249.

14. Ciftci, K., and Levy, R.J. (2001). Enhanced plasmid DNA transfection with lysosomotropic agents in cultured fibroblasts. International journal of pharmaceutics 218, 81–92.

15. Cossart, P., and Helenius, A. (2014). Endocytosis of viruses and bacteria. Cold Spring Harb Perspect Biol 6.

16. Costa, T.R., Felisberto-Rodrigues, C., Meir, A., Prevost, M.S., Redzej, A., Trokter, M., and Waksman, G. (2015). Secretion systems in Gram-negative bacteria: structural and mechanistic insights. Nat Rev Microbiol 13, 343–359.

17. Cullis, P.R., and Hope, M.J. (2017). Lipid Nanoparticle Systems for Enabling Gene Therapies. Mol Ther 25, 1467–1475.

18. de Beer, M., Roverts, R., Heiligenstein, X., Lamers, E., Sommerdijk, N., and Akiva, A. (2021). Visualizing Biological Tissues: A Multiscale Workflow from Live Imaging to 3D Cryo-CLEM. Microscopy and Microanalysis 27, 11–12.

19. Dell’Era, S., Buchrieser, C., Couve, E., Schnell, B., Briers, Y., Schuppler, M., and Loessner, M.J. (2009). *Listeria monocytogenes* L-forms respond to cell wall deficiency by modifying gene expression and the mode of division. Mol Microbiol 73, 306–322.

20. Dubnau, D. (1991). Genetic competence in Bacillus subtilis. Microbiological reviews 55, 395–424.

21. Elkin, S.R., Lakoduk, A.M., and Schmid, S.L. (2016). Endocytic pathways and endosomal trafficking: a primer. Wien Med Wochenschr 166, 196–204.

22. Ellison, C.K., Dalia, T.N., Vidal Ceballos, A., Wang, J.C., Biais, N., Brun, Y.V., and Dalia, A.B. (2018). Retraction of DNA-bound type IV competence pili initiates DNA uptake during natural transformation in Vibrio cholerae. Nat Microbiol 3, 773–780.

23. Errington, J. (2013). L-form bacteria, cell walls and the origins of life. Open Biol 3, 120143.

24. Errington, J., Mickiewicz, K., Kawai, Y., and Wu, L.J. (2016). L-form bacteria, chronic diseases and the origins of life. Phil Trans R Soc B 371, 20150494.

25. Evers, M.J.W., Kulkarni, J.A., van der Meel, R., Cullis, P.R., Vader, P., and Schiffelers, R.M. (2018). State-of-the-Art Design and Rapid-Mixing Production Techniques of Lipid Nanoparticles for Nucleic Acid Delivery. Small Methods 2.

26. Forgac, M. (2007). Vacuolar ATPases: rotary proton pumps in physiology and pathophysiology. Nat Rev Mol Cell Biol 8, 917–929.

27. Forster, B.M., and Marquis, H. (2012). Protein transport across the cell wall of monoderm Gram-positive bacteria. Mol Microbiol 84, 405–413.

28. Friedrich, A., Hartsch, T., and Averhoff, B. (2001). Natural transformation in mesophilic and thermophilic bacteria: identification and characterization of novel, closely related competence genes in Acinetobacter sp. strain BD413 and Thermus thermophilus HB27. Applied and Environmental Microbiology 67, 3140–3148.

29. Gilbreath, J.J., Cody, W.L., Merrell, D.S., and Hendrixson, D.R. (2011). Change is good: variations in common biological mechanisms in the epsilonproteobacterial genera Campylobacter and Helicobacter. Microbiol Mol Biol Rev 75, 84–132.

30. Gregory, M.A., Till, R., and Smith, M.C.M. (2003). Integration site for *Streptomyces* phage phiBT1 and development of site-specific integrating vectors. J Bacteriol 185, 5320–5323.

31. Hahn, J., Albano, M., and Dubnau, D. (1987). Isolation and characterization of Tn917lac-generated competence mutants of Bacillus subtilis. J Bacteriol 169, 3104–3109.

32. Hammond, L.R., White, M.L., and Eswara, P.J. (2019). ¡vIVA la DivIVA! J Bacteriol 201.

33. Hamoen, L.W., Venema, G., and Kuipers, O.P. (2003). Controlling competence in Bacillus subtilis: shared use of regulators. Microbiology (Reading) 149, 9–17.

34. Han, J., Shi, W., Xu, X., Wang, S., Zhang, S., He, L., Sun, X., and Zhang, Y. (2015). Conditions and mutations affecting *Staphylococcus aureus* L-form formation. Microbiology 161, 57–66.

35. Hensel, M., Achmus, H., DeckersHebestreit, G., and Altendorf, K. (1996). The ATP synthase of Streptomyces lividans: Characterization and purification of the F1Fo complex. Bba-Bioenergetics 1274, 101–108.

36. Hoffmann, J., and Mendgen, K. (1998). Endocytosis and membrane turnover in the germ tube of uromyces fabae. Fungal Genet Biol 24, 77–85.

37. Hou, X.C., Zaks, T., Langer, R., and Dong, Y.Z. (2021). Lipid nanoparticles for mRNA delivery. Nat Rev Mater 6, 1078–1094.

38. Inamine, G.S., and Dubnau, D. (1995). ComEA, a Bacillus subtilis integral membrane protein required for genetic transformation, is needed for both DNA binding and transport. J Bacteriol 177, 3045–3051.

39. Jurasek, M., Flardh, K., and Vacha, R. (2020). Effect of membrane composition on DivIVA-membrane interaction. Biochim Biophys Acta Biomembr 1862, 183144.

40. Kieser, T., Bibb, M.J., Buttner, M.J., Chater, K.F., and Hopwood, D.A. (2000). Practical Streptomyces genetics (Norwich: The John Innes Foundation).

41. Kotnik, T. (2013). Lightning-triggered electroporation and electrofusion as possible contributors to natural horizontal gene transfer. Phys Life Rev 10, 351–370.

42. Kruger, N.J., and Stingl, K. (2011). Two steps away from novelty--principles of bacterial DNA uptake. Mol Microbiol 80, 860–867.

43. Kulkarni, J.A., Witzigmann, D., Chen, S., Cullis, P.R., and van der Meel, R. (2019). Lipid Nanoparticle Technology for Clinical Translation of siRNA Therapeutics. Accounts Chem Res 52, 2435–2444.

44. Lande, M.B., Donovan, J.M., and Zeidel, M.L. (1995). The relationship between membrane fluidity and permeabilities to water, solutes, ammonia, and protons. J Gen Physiol 106, 67–84.

45. Lenaz, G. (1987). Lipid fluidity and membrane protein dynamics. Biosci Rep 7, 823–837.

46. Li, L., Wan, T., Wan, M., Liu, B., Cheng, R., and Zhang, R. (2015). The effect of the size of fluorescent dextran on its endocytic pathway. Cell Biol Int 39, 531–539.

47. Liang, W., and W. Lam, J.K. (2012). Endosomal Escape Pathways for Non-Viral Nucleic Acid Delivery Systems. In Molecular Regulation of Endocytosis.

48. MacNeil, D.J., Gewain, K.M., Ruby, C.L., Dezeny, G., Gibbons, P.H., and MacNeil, T. (1992). Analysis of *Streptomyces avermitilis* genes required for avermectin biosynthesis utilizing a novel integration vector. Gene 111, 61–68.

49. Mercier, R., Kawai, Y., and Errington, J. (2013). Excess membrane synthesis drives a primitive mode of cell proliferation. Cell 152, 997–1007.

50. Mulkidjanian, A.Y., Makarova, K.S., Galperin, M.Y., and Koonin, E.V. (2007). Inventing the dynamo machine: the evolution of the F-type and V-type ATPases. Nat Rev Microbiol 5, 892–899.

51. Munch, B., Trtik, P., Marone, F., and Stampanoni, M. (2009). Stripe and ring artifact removal with combined wavelet - Fourier filtering. Opt Express 17, 8567–8591.

52. Muschiol, S., Balaban, M., Normark, S., and Henriques-Normark, B. (2015). Uptake of extracellular DNA: competence induced pili in natural transformation of *Streptococcus pneumoniae*. Bioessays 37, 426–435.

53. Nishida, H. (2020). Factors That Affect the Enlargement of Bacterial Protoplasts and Spheroplasts. Int J Mol Sci 21.

54. Oparka, K.J., Wright, K.M., Murant, E.A., and Allan, E.J. (1993). Fluid-Phase Endocytosis - Do Plants Need It. J Exp Bot 44, 247–255.

55. Patel, S., Kim, J., Herrera, M., Mukherjee, A., Kabanov, A.V., and Sahay, G. (2019). Brief update on endocytosis of nanomedicines. Adv Drug Deliv Rev 144, 90–111.

56. Perona, P., and Malik, J. (1990). Scale-Space and Edge-Detection Using Anisotropic Diffusion. Ieee T Pattern Anal 12, 629–639.

57. Ramijan, K., Ultee, E., Willemse, J., Zhang, Z., Wondergem, J.A.J., van der Meij, A., Heinrich, D., Briegel, A., van Wezel, G.P., and Claessen, D. (2018). Stress-induced formation of cell wall-deficient cells in filamentous actinomycetes. Nat Commun 9, 5164.

58. Ramijan, K., van Wezel, G.P., and Claessen, D. (2017). Genome sequence of the filamentous actinomycete *Kitasatospora viridifaciens*. Genome Announc 5, e01560–01516.

59. Roberts, J., and Park, J.S. (2004). Mfd, the bacterial transcription repair coupling factor: translocation, repair and termination. Current Opinion in Microbiology 7, 120–125.

60. Sato, K., Nagai, J., Mitsui, N., Ryoko, Y., and Takano, M. (2009). Effects of endocytosis inhibitors on internalization of human IgG by Caco-2 human intestinal epithelial cells. Life Sci 85, 800–807.

61. Scheinpflug, K., Krylova, O., and Strahl, H. (2017). Measurement of Cell Membrane Fluidity by Laurdan GP: Fluorescence Spectroscopy and Microscopy. Methods Mol Biol 1520, 159–174.

62. Schindelin, J., Arganda-Carreras, I., Frise, E., Kaynig, V., Longair, M., Pietzsch, T., Preibisch, S., Rueden, C., Saalfeld, S., Schmid, B., et al. (2012). Fiji: an open-source platform for biological-image analysis. Nat Methods 9, 676–682.

63. Shimoni, E., and Muller, M. (1998). On optimizing high-pressure freezing: from heat transfer theory to a new microbiopsy device. J Microsc 192, 236–247.

64. Shitut, S., Shen, M.-J., Claushuis, B., Derks, R.J.E., Giera, M., Rozen, D., Claessen, D., and Kros, A. (2021). Generating heterokaryotic cells via bacterial cell-cell fusion. BioRxiv.

65. Spehner, D., Steyer, A.M., Bertinetti, L., Orlov, I., Benoit, L., Pernet-Gallay, K., Schertel, A., and Schultz, P. (2020). Cryo-FIB-SEM as a promising tool for localizing proteins in 3D. J Struct Biol 211, 107528.

66. Studer, D., Michel, M., and Muller, M. (1989). High pressure freezing comes of age. Scanning Microscopy 3, 253–268; discussion 268.

67. Studer, P., Staubli, T., Wieser, N., Wolf, P., Schuppler, M., and Loessner, M.J. (2016). Proliferation of *Listeria monocytogenes* L-form cells by formation of internal and external vesicles. Nat Commun 7, 13631.

68. Stülke, J., Eilers, H., and Schmidl, S.R. (2009). Mycoplasma and spiroplasma. In Encyclopedia of microbiology, M. Schaechter, ed. (Oxford: Elsevier), pp. 208–219.

69. Stuttard, C. (1982). Temperate phages of *Streptomyces venezuelae*: lysogeny and host specificity shown by phages SV1 and SV2. J Gen Microbiol 128, 115–121.

70. Subramanya, S., Hardin, C.F., Steverding, D., and Mensa-Wilmot, K. (2009). Glycosylphosphatidylinositol-specific phospholipase C regulates transferrin endocytosis in the African trypanosome. Biochem J 417, 685–694.

71. Takahashi, S., Mizuma, M., Kami, S., and Nishida, H. (2020). Species-dependent protoplast enlargement involves different types of vacuole generation in bacteria. Sci Rep 10, 8832.

72. Thomas, C.M., and Nielsen, K.M. (2005). Mechanisms of, and barriers to, horizontal gene transfer between bacteria. Nat Rev Microbiol 3, 711–721.

73. Thottacherry, J.J., Sathe, M., Prabhakara, C., and Mayor, S. (2019). Spoiled for Choice: Diverse Endocytic Pathways Function at the Cell Surface. Annu Rev Cell Dev Bi 35, 55–84.

74. Trombone, A.P., Silva, C.L., Lima, K.M., Oliver, C., Jamur, M.C., Prescott, A.R., and Coelho-Castelo, A.A. (2007). Endocytosis of DNA-Hsp65 alters the pH of the late endosome/lysosome and interferes with antigen presentation. PLoS One 2, e923.

75. Varkouhi, A.K., Scholte, M., Storm, G., and Haisma, H.J. (2011). Endosomal escape pathways for delivery of biologicals. J Control Release 151, 220–228.

76. Vidavsky, N., Akiva, A., Kaplan-Ashiri, I., Rechav, K., Addadi, L., Weiner, S., and Schertel, A. (2016). Cryo-FIB-SEM serial milling and block face imaging: Large volume structural analysis of biological tissues preserved close to their native state. J Struct Biol 196, 487–495.

77. Vischer, N. (2016). Using ImageJ to show “Generalized Polarization” (GP).

78. Woese, C. (1998). The universal ancestor. P Natl Acad Sci USA 95, 6854–6859.

79. Woese, C.R. (2000). Interpreting the universal phylogenetic tree. P Natl Acad Sci USA 97, 8392–8396.

80. Wolff, J.A., and Budker, V. (2005). The Mechanism of Naked DNA Uptake and Expression. Adv Genet 54, 3–20.

81. Wolff, J.A., Malone, R.W., Williams, P., Chong, W., Acsadi, G., Jani, A., and Felgner, P.L. (1990). Direct Gene-Transfer into Mouse Muscle Invivo. Science 247, 1465–1468.

82. Wolfgang, M., van Putten, J.P.M., Hayes, S.F., and Koomey, M. (1999). The comP locus of Neisseria gonorrhoeae encodes a type IV prepilin that is dispensable for pilus biogenesis but essential for natural transformation. Molecular Microbiology 31, 1345–1357.

83. Wu, C., van der Heul, H.U., Melnik, A.V., Lubben, J., Dorrestein, P.C., Minnaard, A.J., Choi, Y.H., and van Wezel, G.P. (2019). Lugdunomycin, an Angucycline-Derived Molecule with Unprecedented Chemical Architecture. Angew Chem Int Ed Engl 58, 2809–2814.

84. Yabu, K. (1991). Formation of Vesiculated Large Bodies of Staphylococcus-Aureus L-Form in a Liquid-Medium. Microbiology and Immunology 35, 395–404.

85. Yanisch-Perron, C., Vieira, J., and Messing, J. (1985). Improved M13 phage cloning vectors and host strains: nucleotide sequences of the M13mp18 and pUC19 vectors. Gene 33, 103–119.

86. Yeh, Y.C., Lin, T.L., Chang, K.C., and Wang, J.T. (2003). Characterization of a ComE3 homologue essential for DNA transformation in Helicobacter pylori. Infect Immun 71, 5427–5431.

87. Zacchetti, B., Smits, P., and Claessen, D. (2018). Dynamics of pellet fragmentation and aggregation in liquid-grown cultures of *Streptomyces lividans*. Front Microbiol 9, 943.

88. Zacchetti, B., Willemse, J., Recter, B., van Dissel, D., van Wezel, G.P., Wösten, H.A.B., and Claessen, D. (2016). Aggregation of germlings is a major contributing factor towards mycelial heterogeneity of *Streptomyces*. Sci Rep 6, 27045.

89. Zhang, L., Ramijan, K., Carrion, V.J., van der Aart, L.T., Willemse, J., van Wezel, G.P., and Claessen, D. (2021). An alternative and conserved cell wall enzyme that can substitute for the lipid II synthase MurG. mBio 12.

90. Zuiderveld, K. (1994). Contrast limited adaptive histogram equalization. Graphics gems, 474–485.

