## Supplementary Figures for "DNA uptake by cell wall-deficient bacteria reveals a putative ancient macromolecule uptake mechanism"

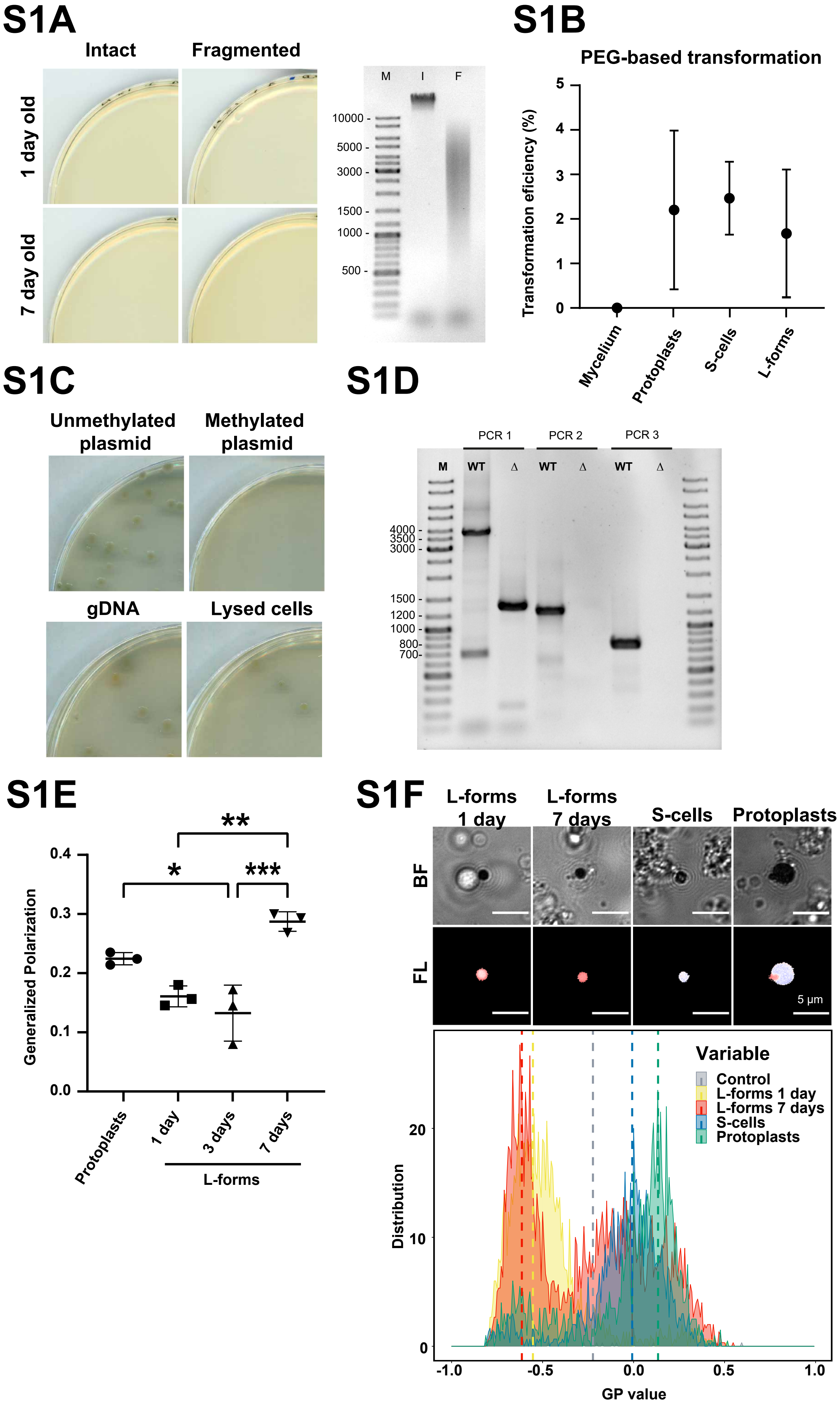

S1A
Intact
Fragmented
1 day old
7 day old
M
I
F
10000 -
5000 -
3000 -
1500 -
1000 -
500 -
S1B
PEG-based transformation
Transformation eficiency (%)
5
4
3
2
1
0
Mycelium
Protoplasts
S-cells
L-forms
S1C
Unmethylated plasmid
Methylated plasmid
gDNA
Lysed cells
S1D
PCR 1
PCR 2
PCR 3
M
WT
Δ
WT
Δ
WT
Δ
4000 -
3500 -
3000 -
1500 -
1200 -
1000 -
800-
700-
S1E
Generalized Polarization
0.4
0.3
0.2
0.1
0.0
**
*
***
Protoplasts
1 day
3 days
7 days
L-forms
S1F
L-forms 1 day
L-forms 7 days
S-cells
Protoplasts
BF
FL
5 µm
Variable
Control
L-forms 1 day
L-forms 7 days
S-cells
Protoplasts
Distribution
20
10
0
-1.0
-0.5
0.0
0.5
1.0
GP value

**Figure S1. Analysis of Natural and Artificial DNA Uptake and Membrane Fluidity of Cell-Wall Deficient Cells and Confirmation of alphaΔcomEA/EC Mutant, Related to Figure 1**

(A) (Left) Transformation plates showing absence of natural transformation upon incubation of 1-and 7-day old L-form alpha with intact or fragmented gDNA of *alphaΔssgB* containing an apramycin resistance cassette. (Right) Gel electrograph of 100 ng intact (I) or fragmented (F) gDNA of *alphaΔssgB* as used in the natural transformation assay.

(B) Polyethylene glycol (PEG)-based transformation efficiency of *K. viridifaciens* mycelium, protoplasts, S-cells and L-forms using plasmid DNA (pRed*) containing an apramycin resistance gene, shown as the percentage of transformed colonies per total colony forming units. Data are represented as mean ± SD, n=3.

(C) PEG-based transformation of alpha using unmethylated or methylated plasmid DNA (pRed*), gDNA or filter-sterilized salt-lysed cells from mutant line *alphaΔssgB*.

(D) Gel electrograph of PCR products from three different PCR mixes to confirm the replacement of *comEA* and *comEC* by an apramycin resistance cassette. WT = gDNA *alpha*; Δ = gDNA *alphaΔcomEA/EC*. Expected products: PCR 1 WT = 3676 bp, mutant = 1294 bp; PCR 2 WT = 1197 bp, mutant = no amplification, PCR 3 WT = 745 bp, mutant = no amplification.

(E) Generalized Polarization (GP) as measure of membrane fluidity of *K. viridifaciens* protoplasts, 1-, 3- and 7- day old L-form *alpha*. Lower GP indicates higher fluidity. *, ** and *** indicate $P \leq 0.05$, 0.01 and 0.001, respectively (one-way ANOVA, $F(3,8) = 19.49$, Tukey post-hoc test, n=3). Data are represented as mean ±SD with individual data points, n=3.

(F) Membrane fluidity of L-form *alpha* (1- and 7-day old), S-cells and protoplasts of *K. viridifaciens*. Top rows show brightfield images and heatmap of fluorescence emission (red to blue colour indicate GP values of -1.0 to 1.0 respectively) of representative cells stained with a Laurdan dye for quantifying the membrane fluidity (BF = brightfield, FL = fluorescence emission). Bottom panel shows frequency distributions of the Generalized Polarization (GP). Lower GP values correspond to higher membrane fluidity indicating that L-forms have more fluid membranes compared to S-cells and protoplasts. Control = cells imaged and analysed without Laurdan staining.

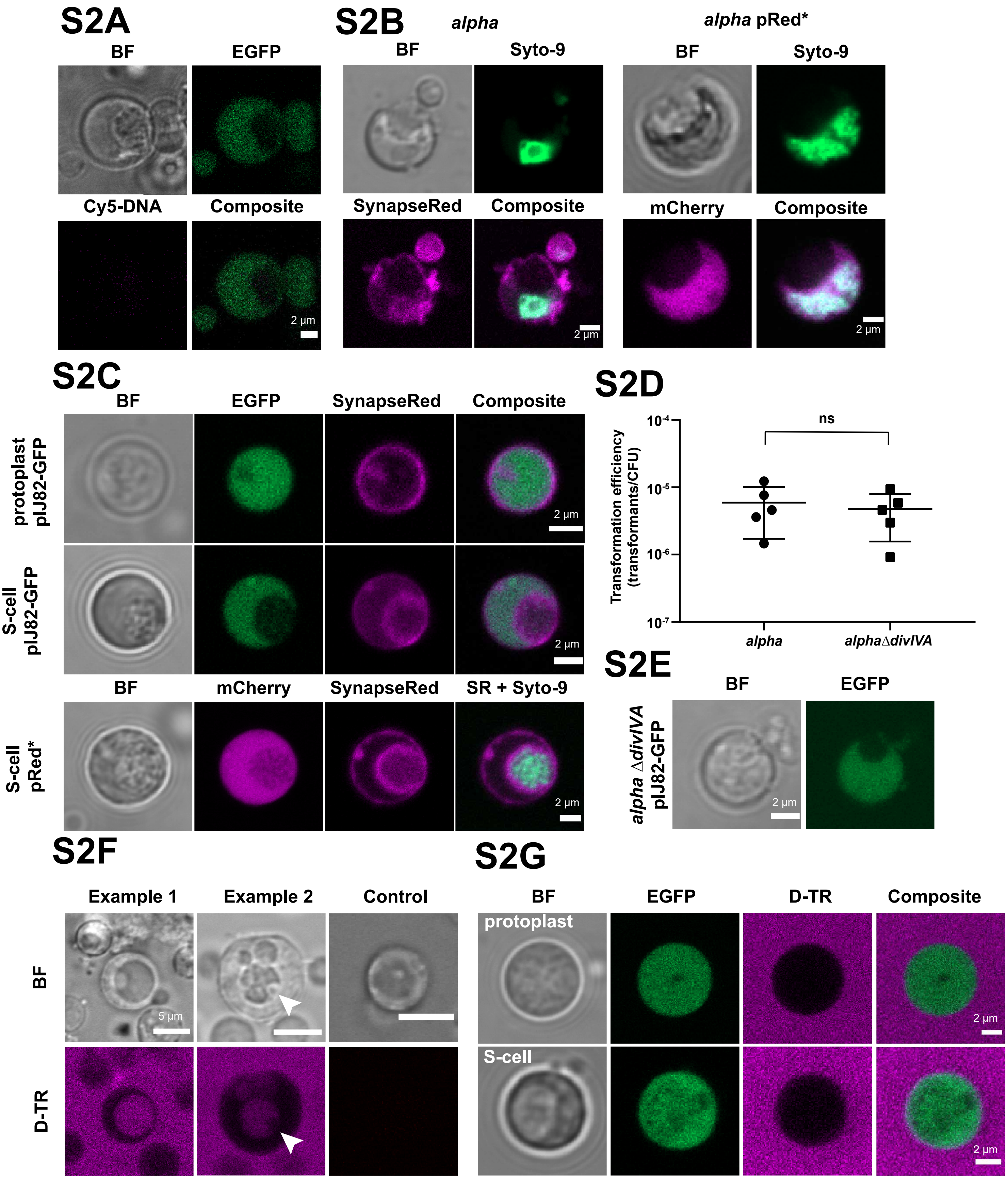
S2A
BF
EGFP
Cy5-DNA
Composite
2 µm
S2B
alpha
alpha pRed*
BF
Syto-9
SynapseRed
Composite
mCherry
2 µm
S2C
BF
EGFP
SynapseRed
Composite
protoplast pIJ82-GFP
S-cell pIJ82-GFP
mCherry
SR + Syto-9
S-cell pRed*
2 µm
S2D
Transformation efficiency (transformants/CFU)
10-4
10-5
10-6
10-7
ns
alpha
alphaΔdivIVA
S2E
alpha ΔdivIVA pIJ82-GFP
BF
EGFP
2 µm
S2F
Example 1
Example 2
Control
BF
D-TR
5 µm
S2G
BF
EGFP
D-TR
Composite
protoplast
S-cell
2 µm

**Figure S2. Analysis of DNA Content, Internal Vesicles and Uptake of D-TR of Cell-Wall Deficient Cells, and effect of divIVA deletion on DNA Uptake, Related to Figure 2**

(A) *alpha* pIJ82-GFP incubated without Cy-5 DNA as fluorescence control.

(B) *alpha* and *alpha* pRed* stained with SYTO-9 (green) to indicate chromosomal DNA. *alpha* is stained with SynapseRed C2M (SynapseRed; magenta) to visualize cell membranes, whereas (absence of) cytosolic mCherry for *alpha* pRed* (magenta) indicates the presence of an internal vesicle.

(C) Protoplasts and S-cells of *K. viridifaciens* pIJ82-GFP producing cytosolic eGFP incubated with SynapseRed for 72 h (top rows), and S-cells of *K. viridifaciens* pRed* producing cytosolic mCherry incubated with SynapseRed (SR) and SYTO-9 for 72 h (bottom row). Chromosomal DNA is visualized using SYTO-9 staining. Note that presence of internal membrane structures causes a reduction in cytosolic fluorescence emission.

(D) Natural transformation assay of 7-day old *alpha* and *alphaΔdivIVA* using pFL-*ssgB*. ns = not significant (two-tailed independent t-test, $t(8)=0.489$, $P=0.638$). Data are represented as mean ± SD with individual data points, n=5.

(E) L-forms without DivIVA can produce internal vesicles as shown for 5-day old *alphaΔdivIVA* pIJ82-GFP producing cytosolic eGFP. Scale bar = 2 μm.

(F) *alpha* incubated with (example 1 and 2) or without (control) Dextran Texas-Red (D-TR; magenta), showing the formation of internal vesicles filled with D-TR. The arrow indicates the presence of a non-fluorescent secondary internal vesicle inside an existing internal vesicle (example 2). Scale bar = 5 μm.

(G) Protoplasts and S-cells of *K. viridifaciens* pIJ82-GFP incubated with D-TR for 72 h. Note that no internalization of D-TR was observed.

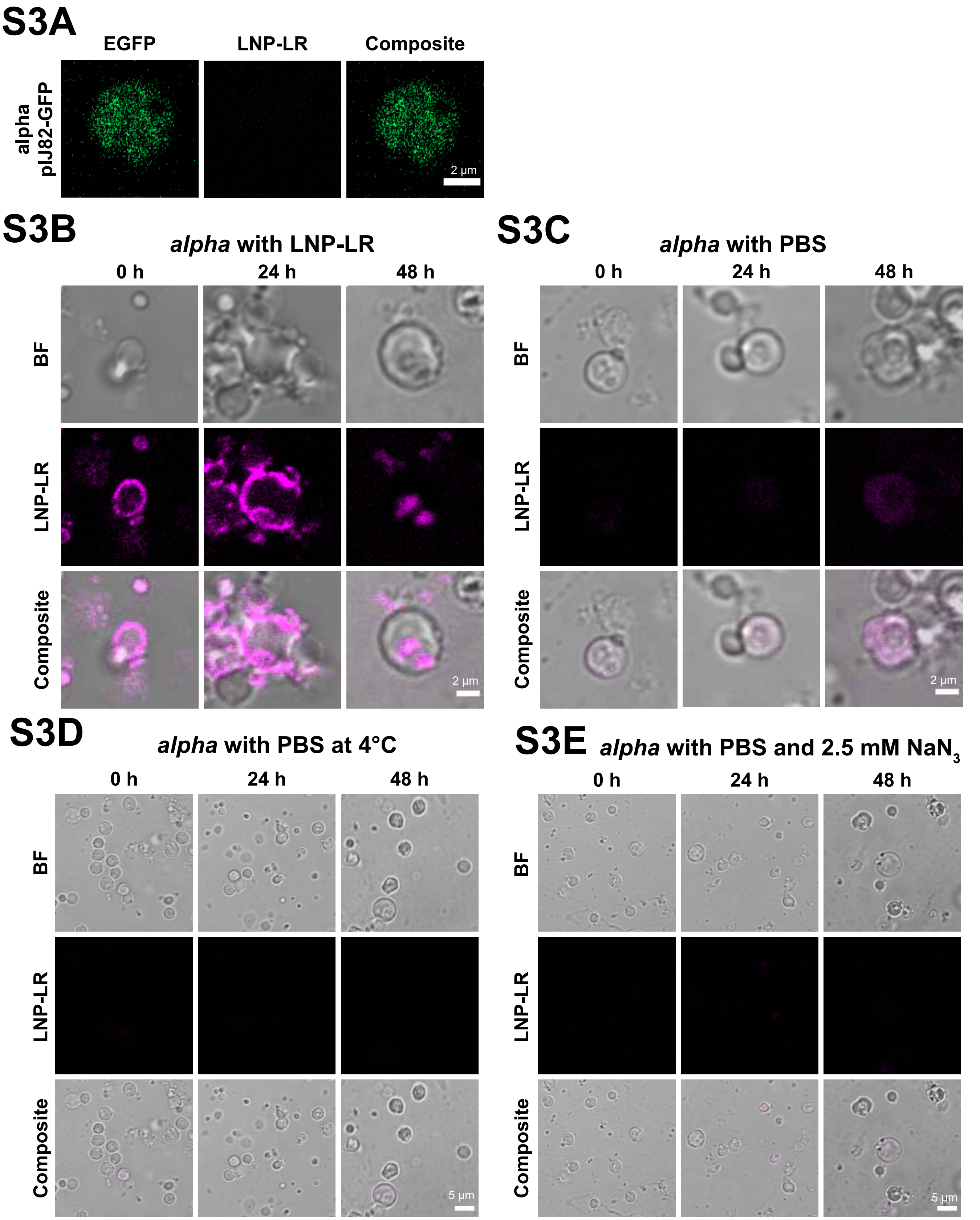


**Figure S3. Uptake of LNP-LR by alpha, Related to Figure 3**

(A) *alpha* pIJ82-GFP incubated without LNP-LR (LNP-Liss Rhod; magenta) as imaging control.

(B-C) *alpha* incubated with (B) or without (C) LNP-LR showing localization of LNP-LR after 0 h, 24 h and 48 h or examples of autofluorescence, respectively.

(D-E) *alpha* incubated with PBS at 4 degrees (D) or with PBS at 30°C in the presence of 2.5 mM sodium azide (E) as control for fluorescence emission. Images were obtained after 0, 24 and 48 h incubation.

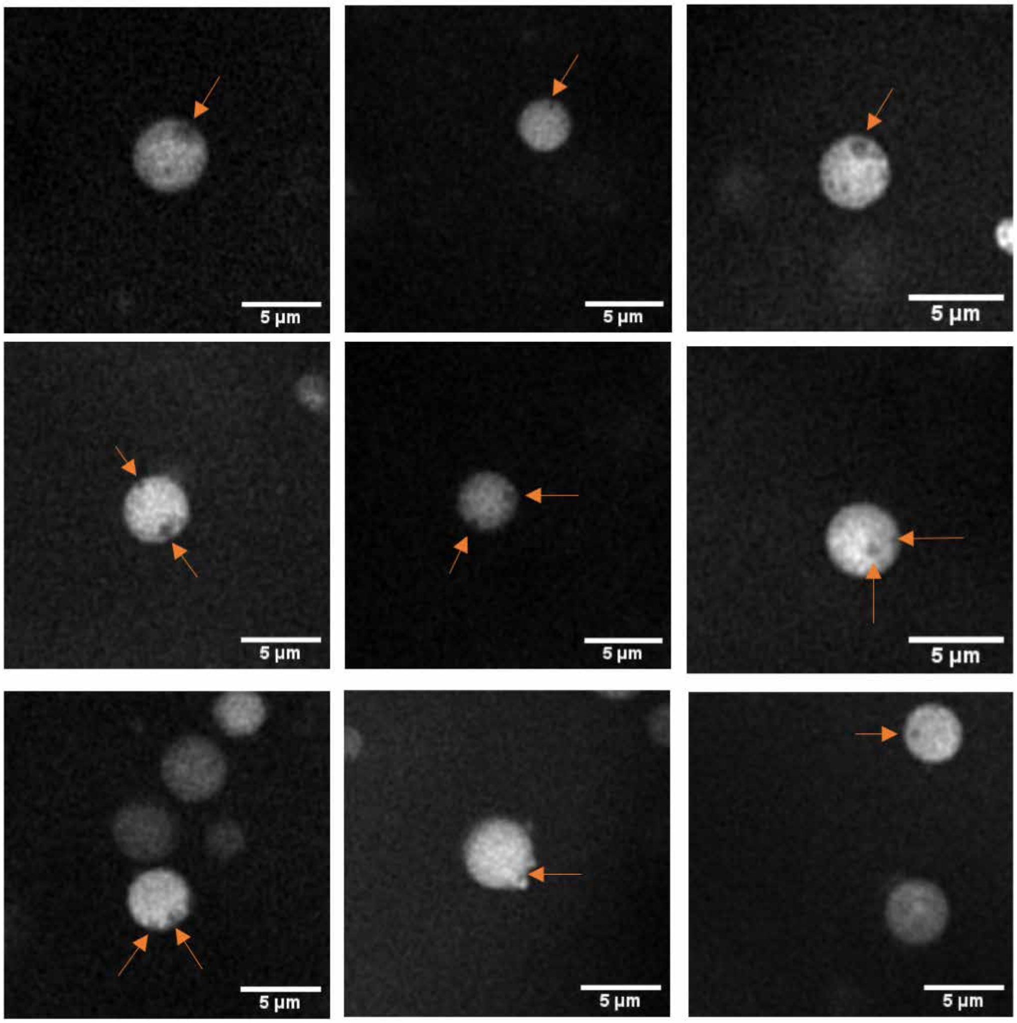


**Figure S4. High Resolution Cryo-Fluorescence of L-forms, Related to Figure 4**
*alpha* pIJ82-GFP imaged using cryo-fluorescence microscopy. Putative vesicles are indicated with arrows. Images were captured using the long distance 100x objective.

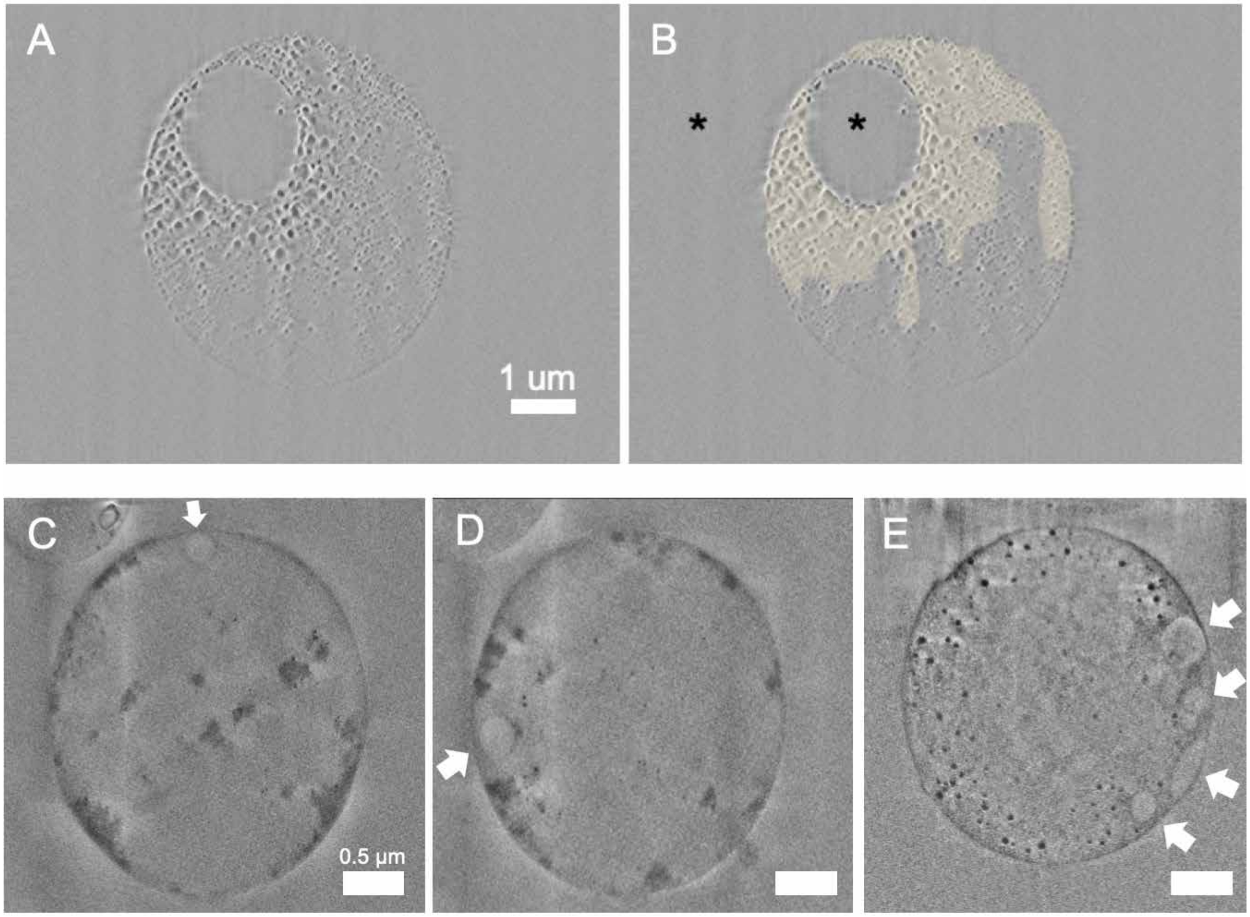


**Figure S5. Over-dose experiment of L-form Cell using FIB-SEM, Related to Figure 4**
(A-B) FIB-SEM slice of over-dose experiment using *alpha* pIJ82-GFP. The yellow colour in B) indicates areas with distinguished beam damage. The vesicle (black asterisk in the center of the cell) seems to be less to none affected by the dose, similar to the medium outside the cell (black asterisk outside of the cell). The image in Figure 4D is taken before this experiment, and Figure 4E is obtained by summing several slices deeper in the cell after acquiring this image.
(C-E) FIB-SEM slices of two cells (C-D correspond to the cell in Figure 4F and E corresponds to the cell in Figure 4 H-K), white arrows indicate vesicles that line the cell membrane. Scale bar in C-E is 0.5 μm.

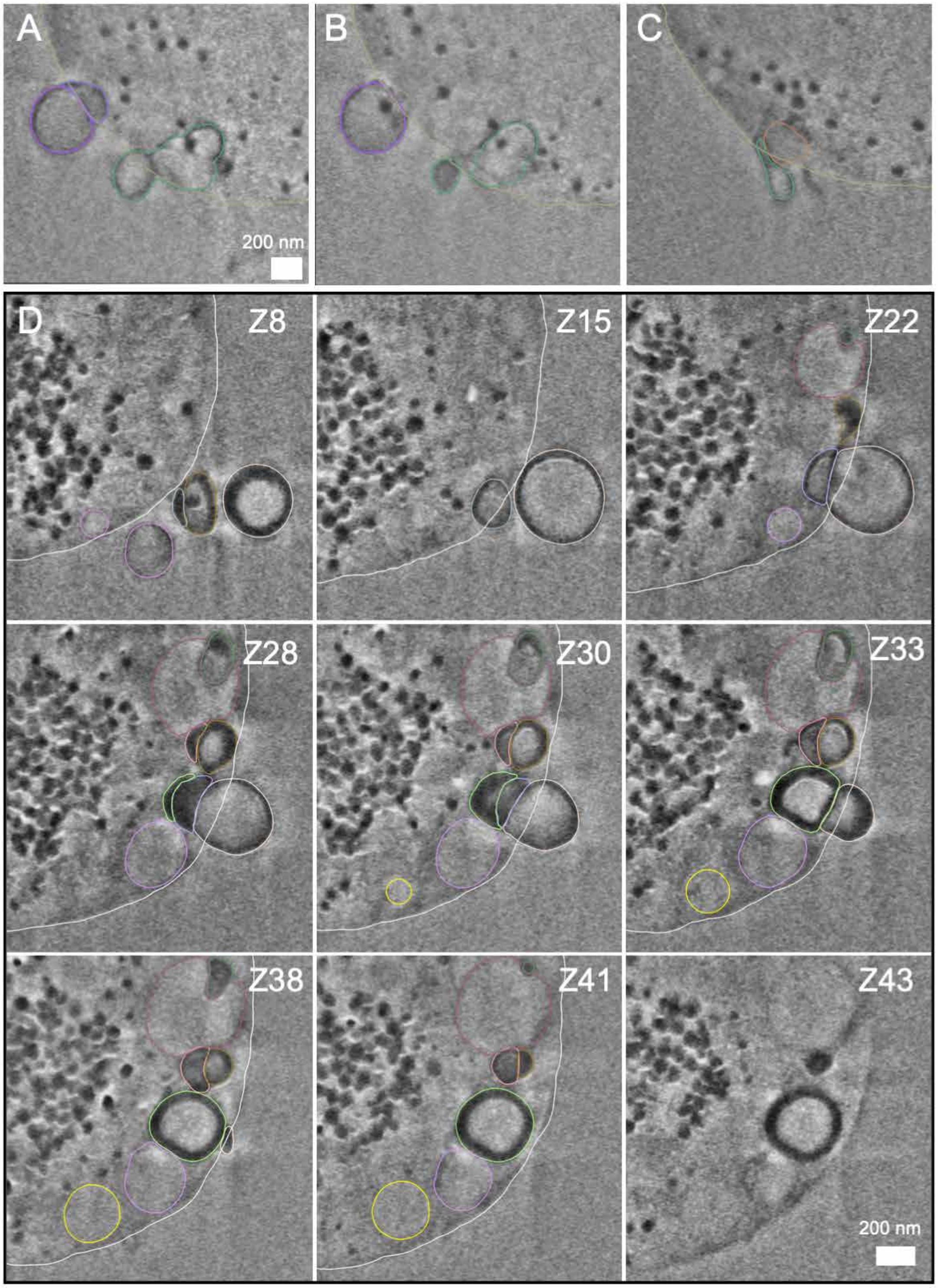


**Figure S6. 3D Segmentation of L-form Vesicles, Related to Figure 4**

(A-C) FIB-SEM slices corresponding to Figure 4I, Jiv and Ki-iii, respectively. Colours correspond to the segmented colours in Figure 4Kiv. Vesicles that are budding out the cells are connected to other vesicles or are elongated inside the cell. Scale bar = 200 nm. See also Video S2.

(D) FIB-SEM slices corresponding to the cell in Figure 4H. Z-number indicates the slice. Colours indicate individual vesicles. See also Video S3.
